# Comparative single-cell transcriptomics of complete insect nervous systems

**DOI:** 10.1101/785931

**Authors:** Benjamin T. Cocanougher, Jason D. Wittenbach, Xi Salina Long, Andrea B. Kohn, Tigran P. Norekian, Jinyao Yan, Jennifer Colonell, Jean-Baptiste Masson, James W. Truman, Albert Cardona, Srinivas C. Turaga, Robert H. Singer, Leonid L. Moroz, Marta Zlatic

## Abstract

Molecular profiles of neurons influence information processing, but bridging the gap between genes, circuits, and behavior has been very difficult. Furthermore, the behavioral state of an animal continuously changes across development and as a result of sensory experience. How behavioral state influences molecular cell state is poorly understood. Here we present a complete atlas of the *Drosophila* larval central nervous system composed of over 200,000 single cells across four developmental stages. We develop *polyseq*, a python package, to perform cell-type analyses. We use single-molecule RNA-FISH to validate our scRNAseq findings. To investigate how internal state affects cell state, we optogentically altered internal state with high-throughput behavior protocols designed to mimic wasp sting and over activation of the memory system. We found nervous system-wide and neuron-specific gene expression changes. This resource is valuable for developmental biology and neuroscience, and it advances our understanding of how genes, neurons, and circuits generate behavior.

## Introduction

Making sense of any complex system involves identifying constituent elements and understanding their individual functions and interactions. Neural circuits are no exception. While recent advances in connectomics (White et al., 1986; Jarrell et al., 2012, Helmstaedter et al., 2013; Takemura et al., 2013; Ohyama et al., 2015; Berck et al., 2016; Eichler et al., 2017; Hildebrand et al., 2017; Eschbach et al., 2019) and live imaging techniques (Ahrens et al., 2013; Prevedel et al., 2014; Chhetri et al., 2015; Lemon et al., 2015; Grimm et al., 2017; Vladimirov et al., 2018) offer unprecedented information about neural connectivity and activity, the task of identifying cell types has traditionally relied on painstaking morphological, functional, or single gene histochemical taxonomy. High-throughput single-cell RNA sequencing (scRNAseq) offers a new way forward by providing a molecular-level identity for each cell via its transcriptomic profile. Importantly, it is also scalable to populations of millions of cells without incurring exorbitant costs. These techniques have already revealed striking heterogeneity in cell populations that is lost in bulk samples. In the fruit fly, efforts are already well underway to produce connectomic (Takemura et al., 2013; Ohyama et al., 2015; Berck et al., 2016; Eichler et al., 2017; Eschbach et al., 2019), activity (Chhetri et al., 2015; Lemon et al., 2015; Grimm et al., 2017; Vladimirov et al., 2018), and behavior atlases (Vogelstein et al., 2014; Robie et al., 2017) of the nervous system. Much work has separately revealed the role that genes (Konopka and Benzer, 1971; Sokolowski 2001) and circuits (Garcia-Campmay et al., 2010, Borst 2014) play in behavior; a major challenge is to combine genes, circuits, and behavior all at once. Single-cell analyses have been performed in parts of adult (Croset et al., 2018; Davie et al., 2018; Konstantinides et al., 2018) *Drosophila* central brain and optic lobe. One study has investigated a small sample of the larval central brain (Alvalos et al., 2019). A comprehensive transcriptomic atlas of the complete central nervous system is the missing piece to the connectivity, activity, and behavior maps that would create the required resource necessary to understand the complex interplay between genes, circuits, and behavior.

To this end, we developed a protocol to capture, sequence, and transcriptionally classify the molecular cell types and cell states of the entire central nervous system of the *Drosophila* larva. We did this across 4 different life stages, providing a developmental profile of gene expression. Given that the *Drosophila* larva has a nervous system of approximately 10,000-15,000 neurons (Hartenstein and Campos-Ortega, 1984; Hartenstein et al., 1987; Truman et al., 1993; Scott et al., 2001), our atlas of 202,107 cells has up to 20X coverage of the entire nervous system and is the largest sequencing effort in *Drosophila* to date. All previously identified cell types were recognizable in our atlas, including motor neurons, Kenyon cells of the mushroom body, insulin-producing cells, brain dopaminergic and serotonergic cells, and all glial subtypes.

While scRNAseq provides nearly complete information about the transcriptional program being used by a cell at the time of collection, a drawback to the technique is a loss of spatial information. We therefore used a recently developed RNA fluorescent *in situ* hybridization (RNA-FISH) protocol to resolve the anatomical location of molecular cell types in the whole larval brain (Long et al., 2017). We combined RNA-FISH with high-resolution Bessel beam structured illumination microscopy to detect and count individual mRNAs within newly identified cells. This technique provides ground truth for the absolute number of a particular RNA in a given molecular cell type at a particular time point. It also provides an opportunity to assess the quantitative capability of our scRNAseq approach.

Larval behavior after hatching is dominated by feeding; when a critical weight is achieved, this behavior switches to “wandering” in preparation for pupation (Bakker et al., 1953). Endocrine and neuroendocrine pathways responsible for this switch have been well characterized (Truman, 2005), but the extent of molecular changes in defined cell types across the nervous system that respond to this neuroendocrine signaling are not known. To investigate such nervous system-wide changes during development, we sequenced the nervous system at four time points in development.

Previous sensory experience alters the behavioral state of an animal. Flies that are hungry form food-associated memories more easily (Krashes et al., 2009) and flies that are intoxicated court more frequently (Lee et al., 2008). Male flies that lose fights are more likely to subsequently exhibit submissive behaviors and to lose second contests while male flies that win exhibit aggressive behavior and are more likely to win later fights (Trannoy et al., 2017). Are internal states controlled transcriptionally at the level of identified cell types and circuits and, if so, how? It is an open question whether memory or internal state will affect gene expression globally or only in restricted cell populations.

In order to discover the nervous system-wide gene expression changes induced by previous experience, we examined gene expression profiles from the nervous systems of animals exposed to two experimental protocols. The first protocol involved presenting repeated pain and fear, by mimicking repeated wasp sting. Fictive stings were induced using optogenetic activation of a small population of well-described interneurons (Ohyama et al., 2015). Of note, no mechanical damage to the animal’s surface occurred with this protocol. The second protocol involved repeated activation of higher-order central brain neurons involved in learning. Using behavioral assays before and after the stimulation, we showed that each of these protocols cause a long-lasting change in the animals’ behavioral state. We then analyzed the effect of fictive sting and repeated activation of the learning center to search for changes in gene expression related to cell state during behavioral learning. We consider these “cell state” genes and find that both entire cell populations and individual neuron types can exhibit cell state changes.

Taken together, these results suggest the powerful role that transcriptomic atlases can play in probing the complex interplay between cell state, circuit function, and behavior.

## Results

### *Polyseq* software performs cell type discovery

A complete transcriptomic atlas of 202,107 single cells from the larval central nervous system was built (Figure 1; Table S1). Nervous systems were captured at four time points in development (1 hour, 24 hours, 48 hours, and 96 hours after larval hatching) and for three nervous system dissections (full CNS, brain only, and ventral nerve cord only). These developmental timepoints and anatomical regions were analyzed separately and in combination. Four non-neuronal and non-glial tissues, including ring gland cells, hemocytes, imaginal disc cells, and salivary gland cells, were also captured and analyzed as outgroups.

**Figure 1.**
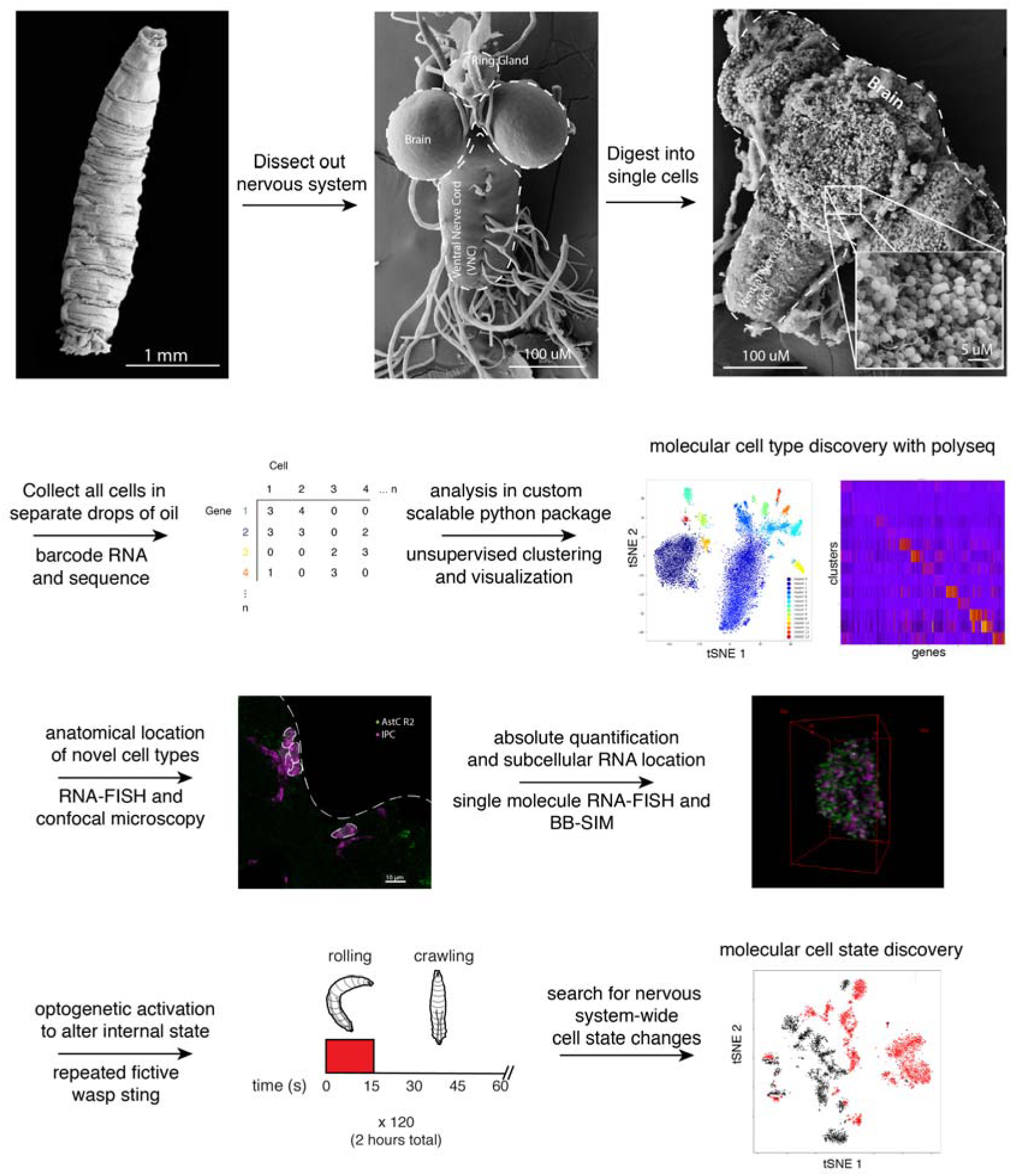
Schematic of full nervous system scRNA-seq collection and analysis. In order to develop a central nervous system-wide transcriptomic atlas of the *Drosophila* nervous system with single-cell resolution, we developed a protocol to digest the entire nervous system into single cells, collected the cells using a microfluidic device (10x Chromium machine, 10x Genomics, Pleasanton, CA), and sequenced the mRNA from each cell. After barcoding and sequencing, a cell by gene matrix is generated. This cell by gene matrix was then analyzed with *polyseq*, a custom python package. In total, 202,107 neurons and glia were sequenced. The anatomical location of newly defined molecular cell types were validated and identified using RNA-FISH with confocal imaging. To push the technique forward, RNA-FISH combined with Bessel beam selective plane illumination microscope (BB-SIM) was used to obtain the absolute quantification and subcellular location of transcripts in these new cell types. Optogenetic manipulations were performed to alter the internal state of the animal, either with two hours of fictive wasp sting or two hours of overactivation of 10% of brain neurons, and scRNAseq was used once more to search for a change in molecular cell state between conditions.

We developed *polyseq* (github.com/jwittenbach/polyseq), an open source Python package, to perform cell type analyses. *polyseq* performs functions of many popular R packages such as *Seurat* or *Monocle* (Trapnell et al., 2014; Satija et al., 2015; Qui et al., 2017; Cao et al., 2019) with significantly improved runtime and the extensibility and modularity of Python. *polyseq* starts with a gene by cell matrix, which is then filtered for high quality cells, normalized, regressed, reduced, and clustered. Visualization can then be performed with tSNE or umap for the full dataset (Maaten and Hinton, 2008; McInnis et al., 2018). The software also includes inbuilt functionality for violin plots and heatmaps. The data remain in a form that is easy to integrate with the vast community of Python packages for further visualization and analysis (full details and analysis examples on github: github.com/jwittenbach/polyseq).

We first used *polyseq* to discover cell type clusters and confirmed that our findings were in agreement with current state of the art analysis methods (Figure 2). In two separate early third instar samples, we found the same cell types when analyzing the data in *Seurat*, *Monocle*, and *polyseq* (Figure 2A). In these samples, the cells separated into seven groups of developing neurons (which included subtypes of adult developing neurons, neuroblasts, and ganglion mother cells), four groups of glia, immune cells, and three groups of larval functional neurons (including distinct motor neuron and Kenyon cell groupings).

**Figure 2.**
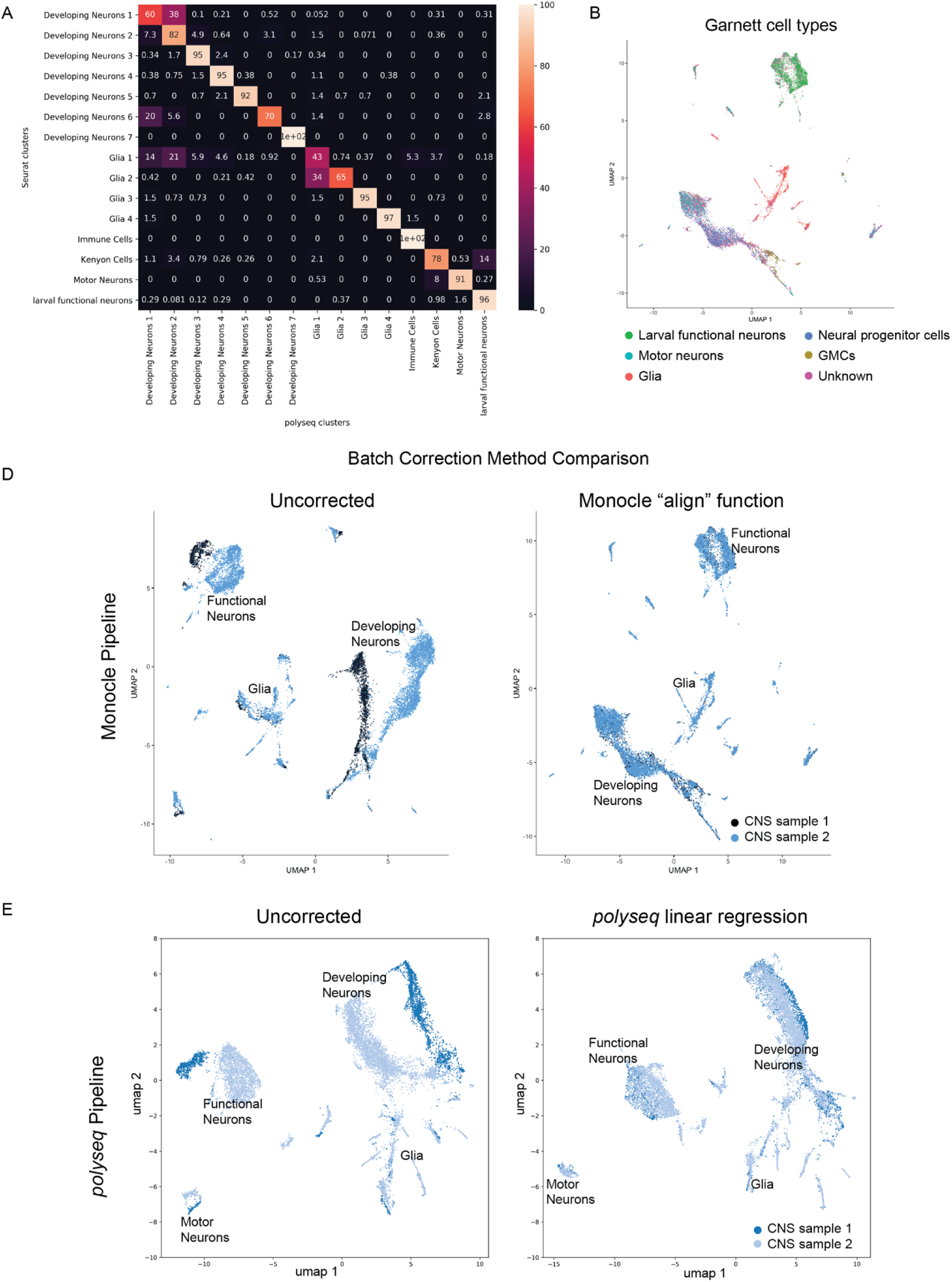
*Polyseq* python package performs cell type discovery and batch correction. A. The same dataset was analyzed in *polyseq* and *Seurat*. A confusion matrix was generated to compare how cells were clustered together. Fifteen clusters were found in both analyses, with 8 of 15 clusters containing more than 90% of the same cells. Clusters that disagreed about cell placement were often differing between two similar clusters (i.e. deciding whether a cell belonged to glia 1 vs glia 2). B. Garnett, a newly developed unsupervised technique, was used to label cell groups with known markers (Pliner et al., 2019). This analysis correctly annotated larval functional neurons, motor neurons, glia, NPCs, and GMCs. It also provides an “unknown” label for cells with low confidence. D,E. Batch correction performance was compared in *Monocle* and *polyseq*. In the plots, cells are separated by a signature related to small differences in sample collection rather than cell type signatures. Monocle’s align function correctly collapses the separated developing neurons (blue and black in “uncorrected” plot) into a comingled group. The linear regression method we implemented in *polyseq* also collapses the sample separations (such as the separation of motor neurons, functional neurons and developing neurons) in the “uncorrected” plot into a single, mixed group in the umap plot following linear regression.

To correct for batch effects, both the align function used in the monocle R package (Haghverdi et al., 2018) and our own linear regression method in *polyseq* were tested (Figure 2B,C). Both methods removed the visible batch effects in the umap plots (i.e., clusters that were made entirely of a single sample due to signal from separate batches collapsed into a single co-mingled population). As additional validation of cell type discovery, we used Garnett, a newly developed machine learning software package in R, to build a classifier based on cell type markers (Pliner et al., 2019). We found consistent results with our own annotation of known and newly discovered markers for larval functional neurons, neural stem cells, motor neurons, kenyon cells, and glia (Figure 2D,E). Using more specific markers for known cells which are small in overall number (such as insulin-producing cells, dopaminergic cells, octopaminergic cells, etc.) led to overfitting of the data. These known gene markers could be used to extract cells of interest without unsupervised methods.

### Developmental profile of gene expression across four life stages

Given that the analysis in *polyseq* met the standards of current state of the art methods, we moved forward with an analysis of developmental timepoints. We built atlases of the entire nervous system at 1 hour, 24 hours, 48 hours, and 96 hours. Within these atlases, it was clear that during development, the cellular composition of the nervous system changes (Figure S1). At 1 hour, the nervous system is primarily larval functional neurons. As development proceeds, the absolute number of larval functional neurons remains relatively constant while the proportion of developing neurons greatly expands.

Having identified the main classes of cells, we investigated developmental trajectories of 12,448 neural progenitor cells (NPCs) (Figure 3). We extracted and combined NPCs cells from three stages of development (1 hour, 24 hours, and 48 hours) and performed an analysis in Monocle (Trapnell et al., 2014; Qiu et al., 2017; Cao et al., 2019) (Figure 3). Garnett was used to predict cell types (Pliner et al., 2019). Known cell ages were used to anchor a psuedotime analysis, which aligned the data from early to late NPCs. Gene expression in these populations revealed known markers (such as *insensible* (*insb*) in Ganglion Mother Cells) and unexpected markers, including long non-coding RNA (*CR31386* in early NBs). IGF-II mRNA-binding protein (*Imp*) and Syncrip (*Syp*) form important gradients that mark NB age (Liu et al., 2015). *Imp* levels decrease with age while *Syp* increases with age – young NBs have high levels of *Imp* and low levels of Syp, intermediate NBs have intermediate levels of Imp and Syp, and older NBs have low *Imp* and high *Syp*. These waves are evident in our data and provide an opportunity to investigate further temporal gene expression gradients.

**Figure 3.**
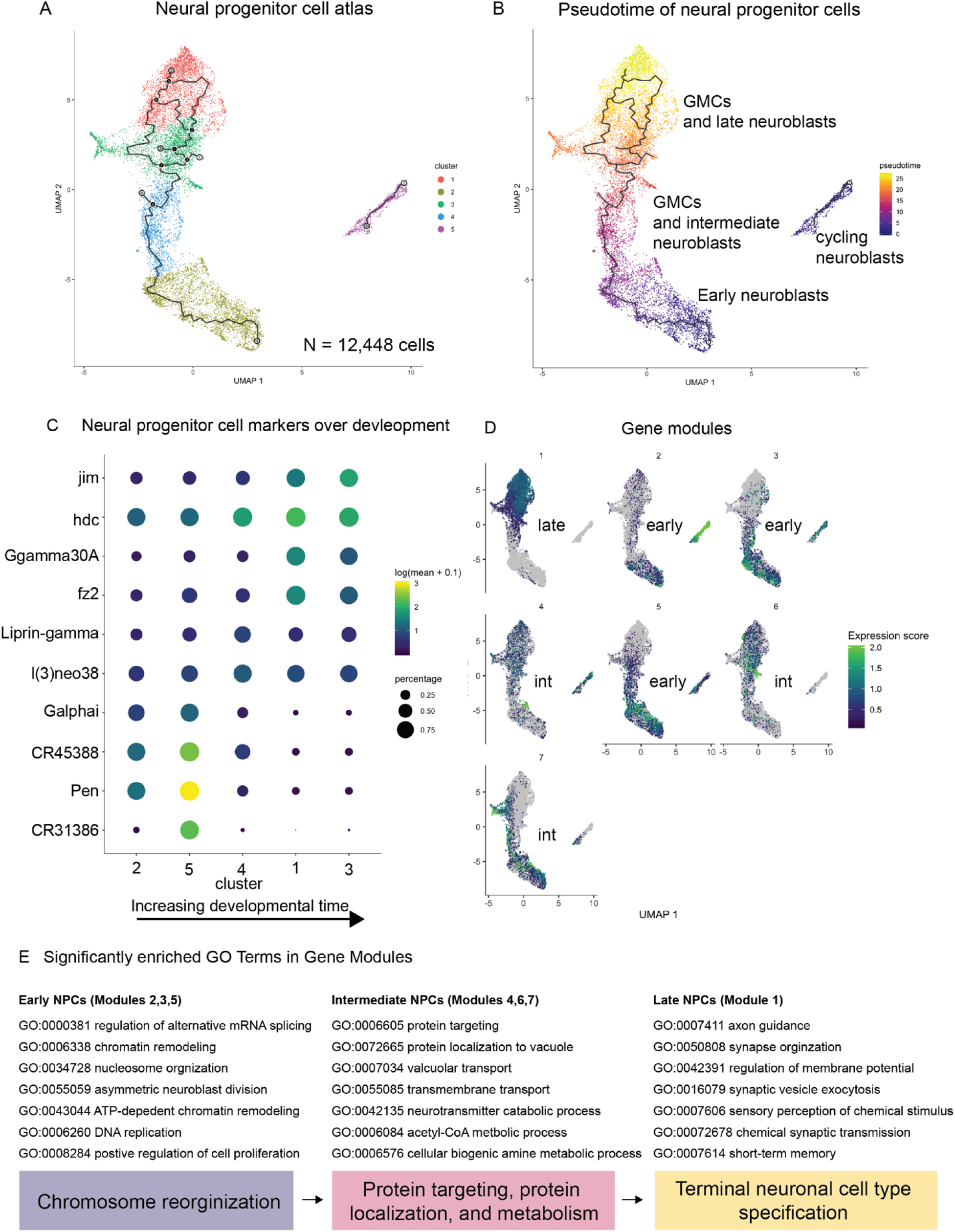
Neural progenitor cell (NPC) atlas reveals gene modules across developmental time. A. A full atlas of first (1H), second (24H), and third (48H) instar larval cells was built, and all NPCs were extracted. These NPCs were then analyzed with Monocle and split into five clusters. B. A pseudotime analysis was performed using known developmental times and separated the data into early, intermediate, and late NPCs. A group of cycling neuroblasts was found to the right of the main NPC dataset in UMAP space. C. Markers for each NPC cluster were extracted and revealed the change in gene expression over developmental time. D. Gene modules were computed and characterized early, intermediate, and late NPCs. As the gene modules represented more developed cells, they were enriched for GO terms (E) which characterized more developed cells (Table S2).

Gene modules were discovered, which characterized populations of early, intermediate, and late NPCs (Figure 3D,E; Table S2). Early NPCs were characterized by the expression of genes important for genome organization, chromatin remodeling, and gene splicing. This allows for a future diversity of cell function and identity. Intermediate NPC gene modules were characterized by genes necessary to build neurons – these included expression of genes important for protein targeting and transport and neurotransmitter synthesis. Late NPCs were enriched for genes which are critical for newly differentiated neurons and circuit construction; significant GO terms included genes required for connecting circuits, such as axon guidance molecules and synapse organization genes, and genes important for circuit function, such as genes involved in memory storage.

### Complete transcriptomic atlas of the larval central nervous system

Next we built an atlas of all cells captured at all stages (Figure 4). Transcriptomic cell types split into seventy clusters (Table S1). These seventy cell types could be grouped into many recognizable groups of cells, including: (1) adult developing brain neurons, (2) adult developing VNC neurons, (3) larval functional brain neurons, (4) larval functional VNC neurons, (5) motor neurons, (6) kenyon cells, (7) brain neuroblasts, (8) VNC neuroblasts, (9) brain ganglion mother cells, (10) VNC ganglion mother cells, (11) glia, (12) hemocytes, (13) imaginal disc cells, (14) salivary gland cells, and (15) ring gland cells.

**Figure 4.**
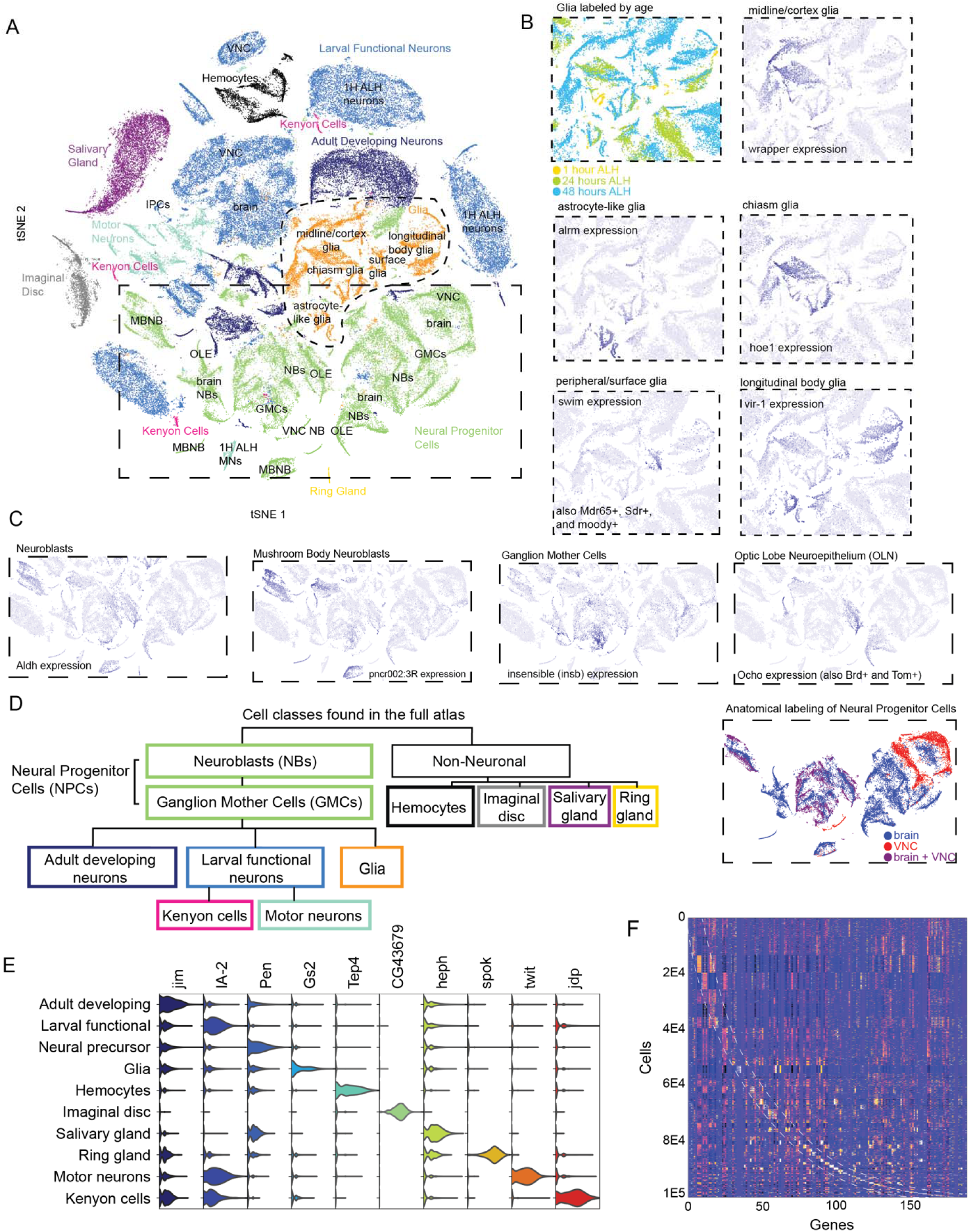
Single-cell transcriptomic atlas of the larval central nervous system. A. t-SNE visualization of high-quality cells colored by cell class. Cells broke into 70 clusters (Figure S4; Table S1) and were post-hoc identified as adult developing neurons, larval functional neurons, neural progenitor cells, glia, hemocytes, imaginal disc, salivary gland, ring gland cells, or subsets of these cells, such as motor neurons, Kenyon cells, neuroblasts, or ganglion mother cells (Table S1). B. Five subtypes of glia, labeled in orange, were found comingled in t-SNE space and could be distinguished by age and function. C. Neural progenitor cells, labeled green in the t-SNE space, split into recognizable classes, including neuroblasts, ganglion mother cells, and optic lobe neuroepithelium. D. Diagram of cell classes contained in the atlas (see Figure S1 for more information). E. Genes that define each cell cluster and cell class were discoverable (Table S1). Violin plots show exemplar genes from each cell class. F. Heatmap of all high-quality cells and the top 3 genes that define each cluster.

Larvae spend much of their life feeding and growing. From initial hatching to pupation, larvae grow significantly in length and mass (Truman et al., 2005). During this growth period, the larval nervous system grows and adds developing adult neurons which remain quiescent during larval life but grow and elaborate their axonal and dendritic arbors during pupation into adult functional neurons (Li et al., 2014). In the atlas, we can identify adult developing neurons through high expression neuronal markers (*nSyb*, *elav*) and a lack of synaptic and neurotransmitter genes (e.g., *VChAT*, *VGlut*). Recent work in the first instar brain showed that adult developing neurons (or undifferentiated neurons) express *headcase* (*hdc*) and unkempt (*unk*) (Avalos et al., 2019). We see this expression continues in adult developing neurons at 24 and 48 hours. Furthermore, we find this group is marked by many more genes, including the actin-binding protein *singed* (*sn*), the zinc finger transcription factor *jim*, and the transcriptional repressor *pleiohomeotic* (*pho*).

Larval functional neurons participate in neural circuits which control sensation and behavior. At larval hatching, embryonic neurons are born, and all the neurons necessary for larval life are functional (Truman and Bate, 1988). These neurons will continue to grow and some populations, such as the Kenyon cells of the mushroom body, will add more neurons throughout development. Identifiable cells at the top level include motor neurons, Kenyon cells, excitatory and inhibitory interneurons, monoaminergic neurons, and neuropeptidergic neurons. A unifying feature of these cells includes expression of classical *Drosophila* neuronal markers (nSyb, elav), however, we also find many other genes that mark the larval functional neuron group robustly. These markers include the transmembrane receptor protein tyrosine kinase activator *jelly belly* (*jeb*), the protein tyrosine phosphatase *IA-2*, the ligand gated chloride channel *Resistant to dieldrin* (*Rdl*), and one of the beta subunits of sodium-potassium pump (*nirvana3; nrv3*).

Monoaminergic neurons play a key role in learning in the fly (Schwaerzel et al., 2003; Selcho et al., 2009). A single top-level cluster was identified with the expression of key monoaminergic synthetic enzymes (*Trh, ple*) and transporters (*DAT, SerT*). Subclustering of this top-level cluster revealed three strong groups, corresponding to serotonergic, dopaminergic, and octopaminergic clusters, identifying previously undescribed markers of these populations of cells which separate one monoamine type from another (Figure S7).

*JhI-21*, a solute carrier 7-family amino acid transporter, is an example of a novel marker found here in 5-HT neurons. This gene encodes for a protein necessary for protein nutrition signaling, which was recently described (Ro et al. 2016; Ziegler et al., 2016; Ziegler et al., 2018). However, these reports describe the importance of the JhI-21 protein peripherally, with no mechanism for the transmitting of nutritional information to the nervous system. Here we see that *JhI-21* is expressed in the serotonin neuron itself, suggesting that serotonin neurons act directly as sensors for the amino acid nutritional state.

Neural progenitor cells include neuroblasts (NBs), intermediate neural progenitors, and ganglion mother cells (GMCs) (Doe, 2017). NBs divide asymmetrically in three ways to produce progeny: type 0 NBs divide into one self-renewing NB and one neuron; type 1 NBs divide into one NB and one GMC; type 2 NBs divide into one neuroblast and one intermediate neural progenitor which then itself divides into a GMC (Doe, 2017). Each GMC then divides terminally to form two neurons or one neuron and one glial cell. Precisely timed patterns of temporal transcription factors guide this development. We are able to investigate these patterns over space and time by collecting the brain and VNC separately and collecting multiple stages of larval development (Figure 3). The mushroom body continues to grow and develop during larval life. We were able to identify mushroom body neuroblasts in our dataset, which were found in brain NB clusters and characterized by high expression of the late neuroblast marker *Syp*, genes for cell cycling, including *pendulin* (*Pen*) and cyclin E (*CycE)*, and by the long noncoding RNA *pncr002:3R* (Figure 4).

Five glial subtypes were recognizable in our atlas, including midline/cortex, astrocyte-like, chiasm, peripheral/surface, and longitudinal body glia (Figure 4) (Freeman, 2015). These glia were identified based on the expression of well-characterized markers, such as *wrapper* and *slit* (*sli*) expression in midline/cortex glia, *alrm* expression in astrocyte-like glia, *hoe1* expression in chiasm glia, *swim* in surface glia, and *vir-1* in longitudinal body glia. In addition, we find *CG5955*, which codes for an L-threonine 3-dehydrogenase, is highly expressed and found specifically in all glia other than longitudinal body glia.

Hemocytes, imaginal disc, salivary gland, and ring gland cells were also captured and sequenced. Hemocytes form the immune system in *Drosophila*. Hemocytes expressed serpent (*srp*), the canonical marker of embryonic hemocytes (Fossett and Schulz, 2001). Hemocytes also had a very high expression of neuropeptide-like precursor 2 (*Nplp2*).

Imaginal discs are embryonic tissues that become adult tissues, such as wings and legs, after metamorphosis. We dissected these cells and sequenced them separately. We found high and specific expression of many uncharacterized genes, including *CG43679*, *CG14850*, *CG44956*, and *CG31698*, among others (Table S1).

Unlike the imaginal disc, which had few genes in common with neurons, salivary gland cells, surprisingly, formed a homogenous group characterized by expression of many genes shared with neurons, such as the nucleo-cytoplasmic shuttling protein *hephaestus* (*heph*), the RNA-binding protein *Syncrip* (*Syp*), the cadherin molecule *Shotgun* (*shg*), and the cell adhesion molecule *Fasciclin 3* (*Fas3*). Given the secretory nature of the salivary gland, it would be interesting to further investigate the evolutionary and developmental relationship between the salivary gland and neurons, especially given that in other animals, such as molluscs, salivary gland cells are secretory and have action potentials (Kater et al., 1978a,b).

The ring gland is critical for transitions in development. The ring gland was characterized by expression of the well-described Halloween genes, including members of the cytochrome P450 family required for ecdysteroid biosynthesis, including *phantom* (*phm*), *spook* (*spo*), *spookier* (*spok*), *disembodied* (*dib*), shadow (*sad*) and *shade* (*shd*) (Gilbert, 2004). The ring gland also has a high expression of the NADP/NADPH phosphatase *curled* (*cu*).

### Validating transcriptomic predictions

Here, we used Insulin-producing cells (IPCs) as illustrative examples to validate scRNA-seq data. IPCs consist of just fourteen neurons in the larval brain (Figure 5A) (Schlegel et al., 2016). These cells participate in circuits which monitor the nutritional status of the larva and function as the larval equivalent of the mammalian pancreas. If IPCs are ablated, larvae and adults are smaller and have a diabetic phenotype, including increased hemolymph trehalose and glucose levels (Rulifson et al., 2002). IPCs secrete insulin-like peptides which regulate hemolymph sugar levels. Graph-based clustering revealed a cluster defined by the strong expression of insulin-like peptide 2 (*Ilp2*) and insulin-like peptide 5 (*Ilp5*) expression, which are canonical markers of IPCs (Figure 5B). By subsetting the data to look only at 96 cells in the putative IPCs cluster, the neurotransmitters and receptors expressed by these could be analyzed (Figure 5C).

**Figure 5.**
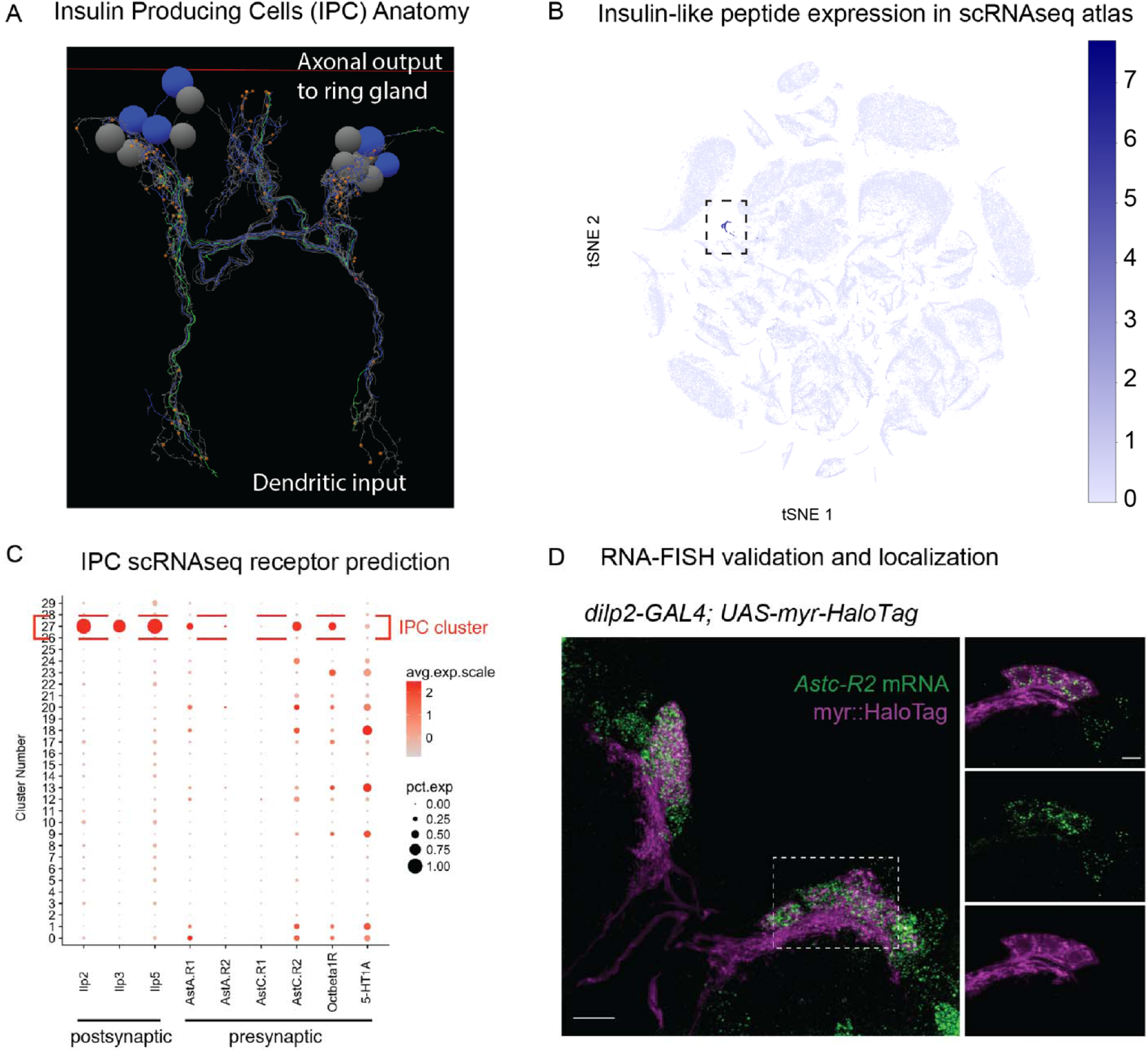
Transcriptomic Atlas predicts previously unknown neuropeptide phenotype for insulin-producing cells and is verified by RNA-FISH. A. Anatomy of insulin-producing cells (IPCs). The IPCs are a group of 7 bilaterally symmetrical neurons which receive input through their dendrites (purple) about the nutritional state of the animal and release insulin-like peptides (ILP2, ILP3, ILP5) through their axons (green), which synapse on the ring gland, to control carbohydrate balance. They are analogous to the vertebrate pancreatic beta islet cells. B,C. The RNAseq atlas built in this study discovered the IPCs as a separate cluster (cluster 27 in C) with expression in the IPCs of receptors for octopamine, serotonin, and allatostatin A, which matched previous literature. Surprisingly, the atlas also suggested the presence of a previously unrecognized receptor in the IPCs for allatostatin C (AstC-R2). D. Detection of AstC-R2 mRNA in IPCs. Maximum-intensity projection of the confocal stack of a brain in which the IPCs are labeled with a fluorescent HaloTag ligand (Magenta) and AstC-R2 mRNA is detected by FISH (green), bar 10µm. Dashed lines outline area where the single z plane is shown on the left panels, bar 5µm. Movie of D is in the supplement (Movie S1).

Previous reports show IPCs are regulated by canonical neurotransmitters. This includes modulation by serotonin through the 5-HT1A receptor and octopamine through the Octbeta1 receptor (Luo et al., 2012) and by the neuropeptide allatostatin A (Hentze et al., 2015). We confirmed this known expression of 5-HT1A and Octbeta1 receptor. In our atlas, we also see the strong expression of additional (previously unknown for these cells) receptors for dopamine (*Dop2R*), glutamate (*GluClalpha*), and Allatostatin C Receptor 2(*AstC-R2*) in IPCs (Figure 5C).

To validate the specificity of our scRNAseq approach for identifying AstC-R2 in ICP cells, we probed AstC-R2 mRNA in a HaloTag reporter line for the ICPs. The overlap between the neurons containing the HaloTag and FISH signals confirmed the sequencing result (Figure 5D). The colocalization of AstC-R2 with 14 IPCs suggests that all ICPs are regulated by AstC through AstC-R2. The discovery of regulation by AstC-R2 updates our model of the regulation of IPCs by adding an additional population of cells that are modulating IPC activity.

### Correlating smRNA-FISH and scRNAseq

In order to determine whether scRNAseq could quantitatively capture the dynamics of expression in a single cell, we compared scRNSeq expression to ground truth expression levels determined using single-molecule RNA-FISH (smRNA-FISH) (Femino et al., 1998). To make analyses more prisise and localized, we compared the RNA levels in a very small population of cells discovered in our atlas that express vesicular glutamate transporter (VGlut) and the neuropeptide Allatostatin C (AstC). We quantified the relative expression of these mRNAs using smFISH and compared the result with scRNAseq (Figure 6).

**Figure 6.**
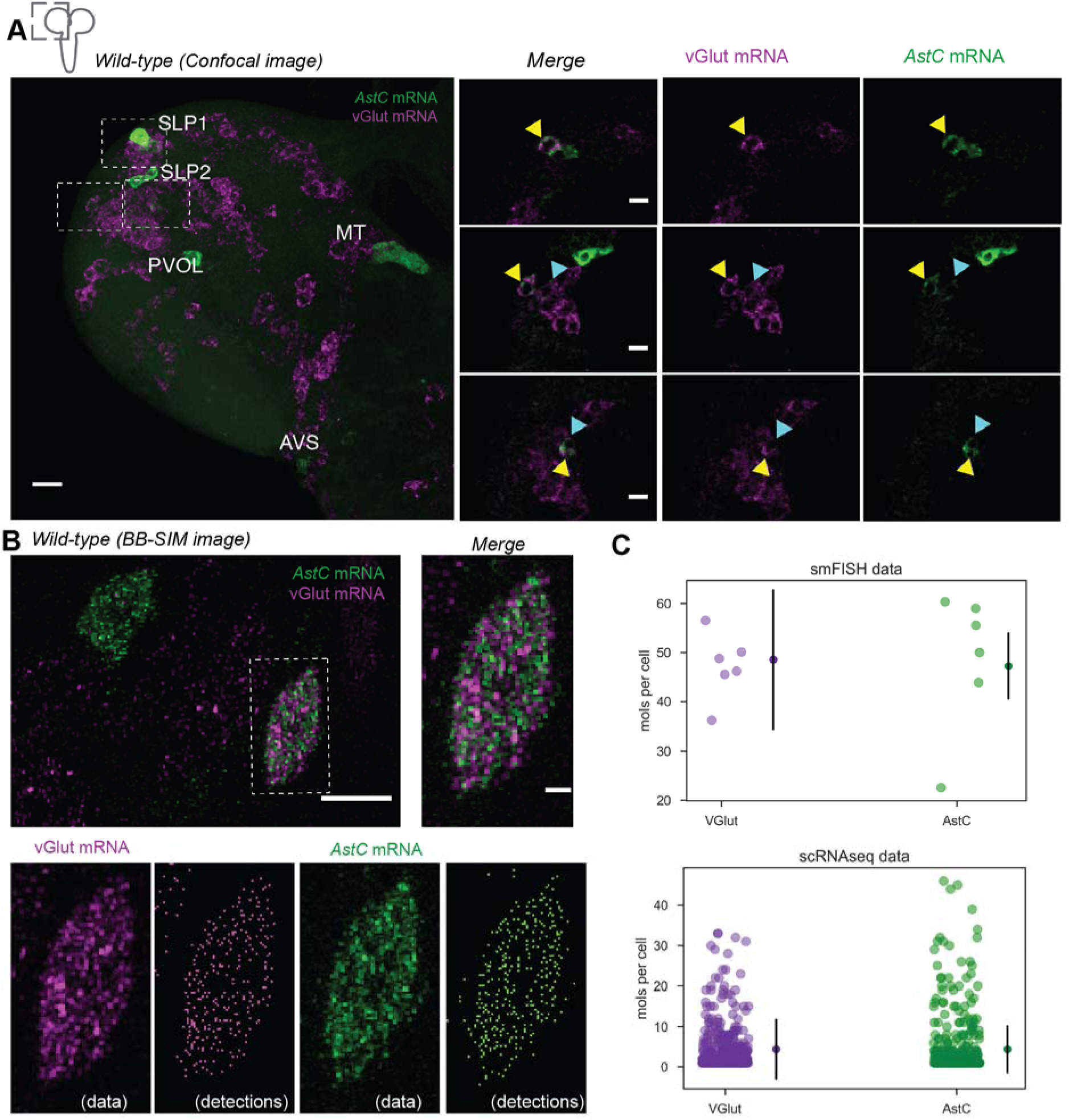
Correlation between single-molecule FISH and scRNAseq. A. Identifying cells that have coexpression of AstC (green) and vGlut (magenta) in the whole brain B. Maximum-intensity projections of BB-SIM stack of the AstC and vGlut mRNA FISH channels, bar 10µm. Dashed lines outline 2 cells that co-express AstC and vGlut mRNAs are shown on the right panels, bar 1µm. Lower panels show individual FISH channel and the reconstructions obtained using the spot-counting algorithm. C. Comparison of the quantification of AstC and vGlut mRNAs between smFISH and scRNAseq.

We used smFISH to probe VGlut and AstC mRNAs simultaneously and obtain quantitative expression levels. We detected 5 groups of cells that contain AstC FISH signals, which was consistent with previous reports of AstC localization (Williamson et al., 2001). We observed 5 pairs of cells that contained both VGlut and AstC. Among the 5 pairs, 1 pair belonged to previously reported SLP1 AstC cells (Figure 6A). We quantified the VGlut and AstC mRNAs in these cells using a Bessel beam selective plane illumination microscope (BB-SIM) (Long et al., 2017). Although quantification of VGlut and AstC mRNAs within individual cell bodies could not be obtained due to the difficulty of segmenting overlapping cell bodies, we were able to obtain an average quantification of VGlut and AstC mRNAs within these 5 pair of cells (Figure 6B). The similarity we obtained for the VGlut and AstC expression ratio between single-molecule FISH and scRNAseq suggested that the relative quantification from scRNAseq was compatible with single-molecule FISH (Figure 6C).

### Optogenetic sting alters expression globally

To investigate if a change in internal state would alter nervous-system-wide gene expression, we examined gene expression profiles from animals exposed to repeated fictive sting. An optogenetic sting was induced by activation of the basin interneurons. The basins are first order interneurons that receive input from nociceptive (pain) and mechanosensory (vibration) sensory neurons (Ohyama et al., 2015). Such optogenetic activation of the brain evokes a rolling escape response (Ohyama et al., 2015), which mimics the natural response to wasp sting or nociceptor activation (Hwang et al., 2007). Of note, no mechanical damage was induced in our protocol.

The basin interneurons were activated for 15 seconds, with a 45 second rest period for a total of 120 activation periods (Figure 7A). Supervised machine learning was used to automatically detect behavior (Jovanic et al., 2017). A rolling escape response was observed at the start of the experiment; by the final activation stimulus, backing up and turning were the predominate responses (Figure 7B).

**Figure 7.**
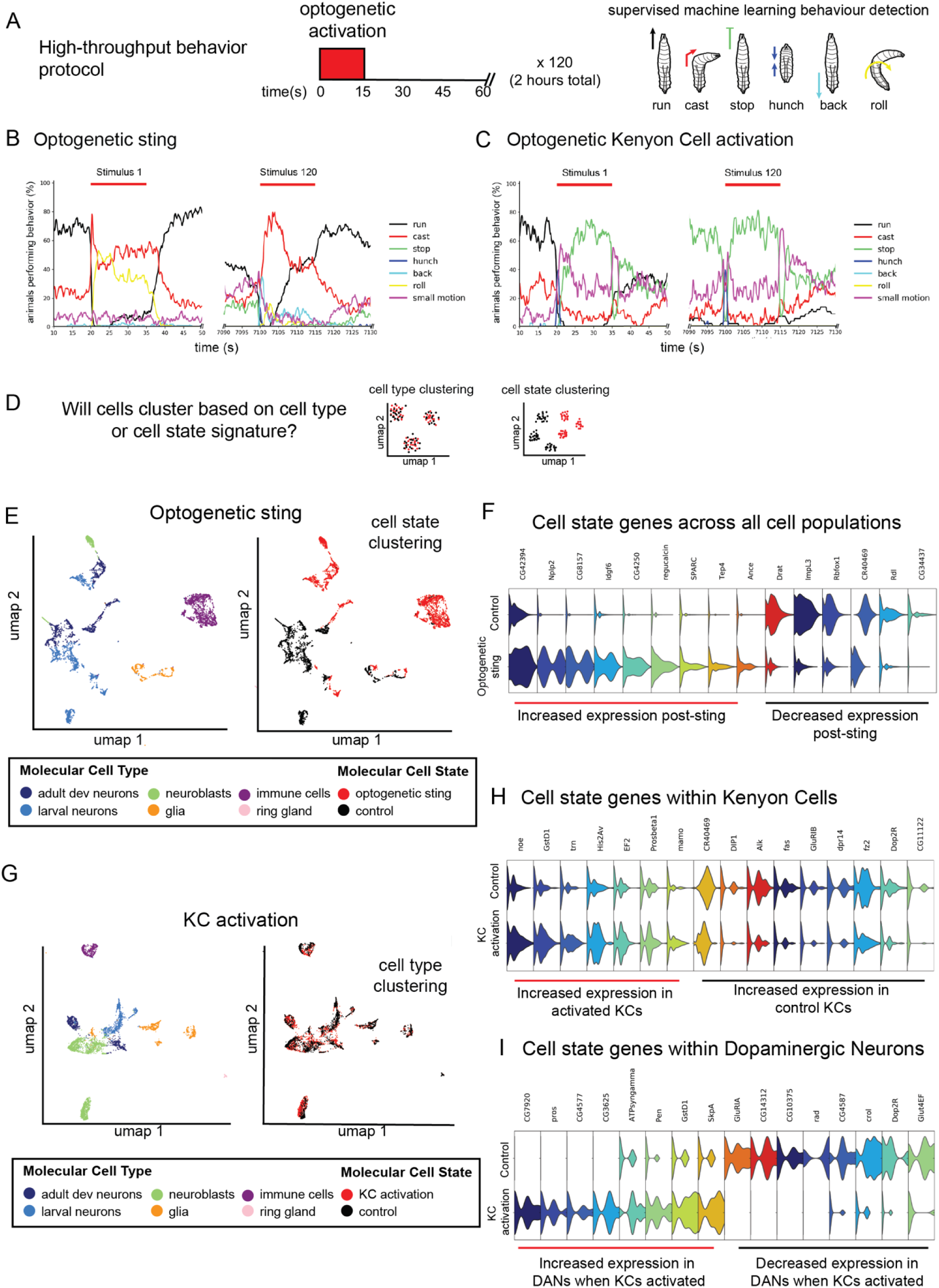
Optogenetic sting and KC activation induce a change in internal state, which is observable as cell state changes in gene expression. A. High-throughput behavior assays were performed using previously described equipment (Ohyama et al., 2013). Fifteen seconds of optogenetic activation were delivered every 60 seconds over a two-hour period. Larval behavior was recorded and supervised machine learning techniques were used to classify behavior as one of six behaviors. B. For the optogenetic sting protocol, animals first respond to basin activation by performing an escape response, including a fast turning response and rolling as previously described (Hwang et al., 2007; Ohyama et al., 2013; Ohyama et al., 2015). Over the course of the experiment, a switch in behavior is observed from rolling to backing up. C. Kenyon cell activation led to a hunch followed by freezing. At the start of training, animals crawled forward to offset. After training, the hunch and freezing response remained at the onset, but the offset response switched to a turn rather than a forward crawl. D. Transcriptomic atlases were produced from animals that underwent each behavior protocol and matched controls. We were primarily interested in whether cell state or cell type clustering would be observed after altering internal state (sting) or overactivating the memory center (KC activation). Cell type clustering would result in cells from activated animals and controls mixing in the same clusters while cell state clustering would lead to separation of cells from activated and control animals. E,F. The optogenetic sting protocol led to cell state clustering. All cell classes were identifiable for both conditions. A large immune cell population was specific to the sting condition, suggesting an immune response to a fictive wasp sting (Figure S8). F. Cell state genes were discovered with upregulated and downregulated expression that separated all cell populations. G-I. KC activation led to cell type clustering and local cell state changes. Even though a larger total number of cells were activated (∼200 KCs versus 64 basins), a less dramatic switch was observed in behavior and in the transcriptome. There were differentially expressed genes in two key populations of cells in the learning and memory center: KCs and dopaminergic neurons.

Control animals of the same age were collected from the same food plate as experimental animals and placed on an agar plate in the dark for two hours. Immediately following the sting protocol, 2-4 animals from each group underwent the scRNAseq protocol. We were primarily interested in searching for cell state genes which could drive cell state clustering (Figure 7D). If cells from experimental animals and controls are analyzed together, will cells cluster together based on cell type (independent of treatment group) or cell state (dependent on treatment group)? If cell type clustering is observed, it suggests that any changes in cell state induced by our protocol are minor compared to cell type-specific features. But if cell state clustering is observed, it is evidence that experience driven changes are at least of comparable importance to cell type in determining genetic cell state.

The optogenetic sting protocol led to cell state clustering. Transcriptomic data from cells isolated from activated and control brains were normalized and analyzed in the same mathematical space, but the clustering that was observed was based on cell state (i.e., clustering was driven by whether the cells came from a “stung” animal or an “unstung” control). Cell state genes that differed between the stung and unstung controls were discovered (Figure 7F).

Cell state genes were most evident in larval functional and developing neurons, including motor neurons, cholinergic, and neuropeptidergic cells. Genes that were upregulated in motor neurons following the sting protocol included non-coding RNA (*CR40469*), carbohydrate metabolic enzymes (lactate dehydrogenase, *ImpL3)*, and the ethanol-induced apoptosis effector, *Drat*.

In addition to cell state genes within the nervous system, a large group of immune cells was sequenced in the sting state, with particularly high expression of neuropeptide-like precursor 2 (*NPLP2*), a gene which has been observed in phagocytic immune cells (Fontana et al., 2012). RNA-FISH in sting and control conditions revealed this transcript is not higher in the neurons but suggested that the lymph gland is being activated and ramping up cell numbers in the sting condition (Figure S8). Strong evolutionary selection pressure exists on the larva to survive predation by parasitic wasps (Kraaijeveld and Godfray, 1997). Larvae can survive by using their immune system to encapsulate and prevent the hatching of internalized parasitic wasp eggs. Previous work has investigated the role of signaling detected by mechanical damage of the sting and the presence of the foreign body (wasp egg) inside the larva (Sorrentino et al., 2002). Our transcriptomic data suggest the immune system may also respond to a currently unknown signal generated directly by the nervous system.

### Learning center overactivation alters expression locally

In a second behavior protocol, we activated all higher-order central brain neurons involved in learning and memory, called Kenyon cells (KCs). Similar to the fictive sting, we observed a change in behavioral response at the start and end of the training. At the start of training, animals hunch and arrest movement at the onset of activation and crawl forward at the offset of activation (Figure 7C). At the end of the training, animals continue to hunch and stop at the onset, but a larger fraction (∼80%) perform a small motion before turning rather than crawling forward to offset. Also, this protocol not only altered animals response to the optogenetic activation of KCs but also drastically altered behavior after activation. Animals greatly increased the probability of stopping and reduced the probability of crawling.

To discover potential molecular changes that could drive these behavioral changes, we analyzed the transcriptomes of animals exposed to these optogenetic training protocols and compared them to controls. Unlike the global changes in gene expression following an optogenetic sting, we detected changes in the transcriptomic state of many fewer cell types following repeated activation of the higher-order brain neurons involved in learning and memory (Figure 7G-I). We discovered a number of interesting candidate genes that were upregulated in an activity-dependent way in Kenyon cells and dopaminergic neurons, which are key cell populations in the learning and memory center (Figure 7;Table S3,S4).

Cell state genes with differential expression between KC activated brains and controls separated local groups of cells within clusters. Changes were observed in KCs and dopaminergic neurons (DANs) (Figure 7H,I). Cell state genes were not limited to previously described activity-related genes. They included long non-coding RNA (*noe*, *CR40469*), chromatin remodeling (*His2AV*, *mamo*), axon guidance (*trn*, *DIP1*, *fas*, *dpr14*, *fz2*), and receptor genes (*Dop2R*).

## Discussion

This work makes several contributions to the field. First, we present the first full transcriptomic atlas of the entire central nervous system at the single-cell resolution. Second, we use super-resolution microscopy to compare single-molecule RNA-FISH with scRNAseq in the *Drosophila* larva. By combining these two techniques – the first providing information about the complete collection of RNA present in a cell and the second providing full anatomical, subcellular, and absolute quantification of a chosen RNA(s) – we provide a resource for the field of *Drosophila* neurobiology and provide an example of complementary methods for building and validating single-cell molecular atlases. Third, we provide an experimental paradigm for discovering a molecular signature of internal state and use this paradigm to uncover drastic gene-expression changes that accompany a state of stress evoked by repeated “optogenetic” predator attack. Our atlas is therefore a powerful resource for developmental biology, neuroscience, and evolutionary biology.

Separating cell type from cell state is a key challenge for transcriptomic cell atlases. In order to understand the changes in a specific cell type between health and disease, for example, it will be necessary to be able to find the same cell type among differing conditions. Here we show that such a distinction can be discovered in cases where molecular cell state is significantly altered. Furthermore, we show that even in a nervous system with drastically different (and unnatural) activity patterns in a sizable population of highly interconnected neurons (here 200 of 10,000 or 2% of the nervous system), limited changes may be observed across the entire nervous system but changes can be observed in specific cell types. We foresee such techniques being useful to investigate a wide range of internal state and cell state changes, from sleep to parasitism to circadian rhythms.

Single-cell transcriptomic atlases are the missing piece required for the combined analysis of genes, circuits, and behavior. Our work here shows that transcriptomic atlases can be reliably built for multiple developmental stages of the *Drosophila* larva. Furthermore, we show that optogenetic manipulations of internal state can alter gene expression in a context-dependent manner. By adding a transcriptomic atlas to the existing atlases of neuron connectivity, neuron activity, and behavior, we have set the stage for a more complete understanding of the principles that underlie the complex interplay of genes, circuits, and behavior.

## Acknowledgments

We thank J. Grimm for sharing reagents; J. Etheredge for fly stocks; S. Harrison, M. Mercer, and the Janelia Fly Core for assistance with fly husbandry; A. Lemire and K. Aswath of Janelia Quantitative Genomics for assistance with sequencing.

## Author contributions

B.T.C., L.L.M., and M.Z. conceptualized the study; B.T.C., J.D.W., X.L., and J.Y. performed all data analysis and visualization; B.T.C., J.D.W., J.-B.M., and J.Y. performed all software development, B.T.C., X.L., J.L., A.B.K., and L.L.M. performed the investigation; B.T.C., J.D.W., and X.L. curated the data; B.T.C., X.L., A.B.K., and L.L.M. developed methodology and performed validation, R.H.S., L.L.M. and M.Z. acquired funding, provided resources, and performed project administration; S.T., R.H.S., L.L.M. and M.Z. supervised the study; B.T.C., J.D.W. and X.L. wrote the original draft; B.T.C., X.L., J.D.W., R.H.S., L.L.M. and M.Z. edited the final manuscript; all authors reviewed the final manuscript.

## METHODS

### CONTACT FOR REAGENT AND RESOURCE SHARING

Further information and requests for resources and reagents should be directed to and will be fulfilled by the Lead Contact, Marta Zlatic.

### EXPERIMENTAL MODEL AND SUBJECT DETAILS

#### Fly stocks

*Drosophila* larvae were grown on standard fly food at 25°C and kept in 12-hour day/night light and dark cycle. Vials were timed by collecting eggs on a new food plate over the course of one hour.

Please see Key Resources Table for *Drosophila* lines used in this study.

### METHOD DETAILS

#### Single cell isolation

*Drosophila* larvae were dissected at 1 hour, 24 hours, 48 hours, or 96 hours after larval hatching (ALH). All dissections were performed in a cold adult hemolymph solution (AHS) with no calcium or magnesium at pH 7.4. Quality of single cell isolation was investigated by visual inspection with compound and confocal microscopy. Samples were placed on ice during waiting periods. Samples were isolated and run on the 10x Chromium Single Cell 3’ immediately after cell dissociation.

First, the complete central nervous system (CNS) was dissected from every animal. The dissected nervous systems were kept in cold AHS on ice. For those samples where the brain and the ventral nerve cord (VNC) were sequenced separately, the separation of the brain from the VNC was performed using fine-tipped forceps and MicroTools (Cat #: 50-905-3315, Electron Microscopy Sciences). The time from digestion (the part of the protocol most likely to induce cell stress) to on the 10x Genomic instrument was never longer than 30 minutes.

After separation of the brain from the VNC, the desired tissue was placed in 18 μL of AHS on ice. Once all samples were prepared, 2 μL of 10x neutral protease (Cat #: LS02100, Worthington Biochemical Corp, Lakewood, NJ, USA) was added to a final volume of 20 μL. The intact brain tissue was digested for 5 minutes. The tissue was then transferred to a fresh drop of 20 μL of AHS.

Each sample was triturated with a clean, thinly pulled glass electrode until no tissue was visible under a dissection scope. All debris (pieces of nerve and undigested neuropile) was removed. Samples with fluorescent markers were observed under a fluorescence microscope to approximate cell density. The samples were then loaded onto the 10x Chromium chip.

#### 10X Genomics

Single cell capture and library construction was performed using the 10x Chromium machine and the Chromium Single Cell 3’ v2 Library and Gel Bead Kit (10x Genomics, Pleasanton, CA). Manufacturer’s recommendations were followed for cell collection and library construction. Libraries were sequenced with an Illumina HiSeq following manufacturer’s instructions.

#### mRNA *in situ* hybridization

FISH probes were designed based on transcript sequences using the online Stellaris Designer and purchased from Biosearch Technologies. Probe sequences for AstC and vGlut were previously reported (Long et al,. 2017; Diaz et al., 2019), and probe sequences for AstC-R2, Hug, NPNL2 are in Table S5. Each probe is 18-22nt long with a 3’ end amine-modified nucleotide that allows directly couple to an NHS-ester dye according to the manufacturer’s instructions (Life Technologies). Dye-labeled probes were separated from the excess free dyes using the Qiagen Nucleotide Removal Columns. FISH protocol was described previously (Long et al,. 2017; Diaz et al., 2019). The brains of 3rd instar larvae were dissected in 1xPBS and fixed in 2% paraformaldehyde diluted PBS at room temperature for 55 min. Brain tissues were washed in 0.5% PBT, dehydrated, and stored in 100% ethanol at 4°C. After exposure to 5% acetic acid at 4 °C for 5 minutes, the tissues were fixed in 2% paraformaldehyde in 1xPBS for 55 min at 25 °C. The tissues were then washed in 1 PBS with 1% of NaBH4 at 4 °C for 30 min. Following a 2 hour incubation in prehybridization buffer (15% formamide, 2 SSC, 0.1% Triton X-100) at 50 °C, the brains were introduced to hybridization buffer (10% formamide, 2 SSC, 5 Denhardt’s solution, 1 mg/ml yeast tRNA, 100 μg/ml, salmon sperm DNA, 0.1% SDS) containing FISH probes at 50 °C for 10 h and then at 37 °C for an additional 10 h. After a series of wash steps, the brains were dehydrated and cleared in xylenes.

#### Confocal and BB-SIM Imaging

For confocal imaging, the tissues were mounted in DPX. Image Z-stacks were collected using an LSM880 confocal microscope fitted with an LD LCI Plan-Apochromat 25x/0.8 oil or Plan-Apochromat 63x/1.4 oil objective after the tissue cured for 24 hours. For single-molecule imaging, we use a previous described Bessel beam selective plane illumination microscope (BB-SIM). Detail construction of microscope and the imaging procedure is described previously (Long et al., 2017). Briefly, this BB-SIM is engineered to image in medium matched to the measured refractive index (RI) of xylene-cleared *Drosophila* tissue with axial resolution of 0.3 µm and lateral resolution of 0.2 µm. For BB-SIM imaging, the tissues were mounted on a 1.5×3mm poly-lysine coated coverslip attached to a 30mm glass rod. The imaging process requires the objectives and tissues immersed in the imaging medium consist with 90% 1,2-dichlorobenzene, 10% 1,2,4-trichlorobenzene with refractive index = 1.5525. Two orthogonally mounted excitation objectives are used to form Bessel beams, which are stepped to create an illumination sheet periodically striped along x or y, while a third objective (optical axis along the z direction) detects fluorescence. To employ structured illumination analysis, we collect multiple images with the illumination stripe pattern shifted to tile the plane in x, and repeat the process orthogonally to tile the plane in y. The sample is then moved in z, and the imaging repeated, and so on to image the 3D volume.

#### High-throughput Automated Optogenetic Behavior Experiments

For the sting mimic experiments, 72F11-GAL4 males were crossed to UAS-CsChrimson virgins (stock information in Key Resources Table). For Kenyon cell overactivation, 201Y-GAL4 males were crossed to UAS-CsChrimson virgins. Larvae were grown in the dark at 25°C. They were raised on standard fly food containing trans-retinal (SIGMA R2500) at a final concentration of 500 µM. Activation was performed in a high-throughput optogenetic behavior rig described previously (Ohyama et al., 2013). About 40 animals were placed in a 25 x 25 cm^2^ dish covered with clear 4% agar.

Neurons were activated using a red LED at 325 µW/cm^2^ illuminated from below the agar dish for 15 seconds with with a 45 second rest period for a total of 120 activation periods (Figure 7A). Supervised machine learning was used to automatically detect behavior (Jovanic et al., 2017). Control animals of the same age were collected from the same food plate as experimental animals and placed on an agar plate in the dark for two hours. Immediately following the sting protocol, 2-4 animals from each group were dissected and cells were collected using the 10X Genomics protocol described above.

### QUANTIFICATION AND STATISTICAL ANALYSIS

#### scRNA-seq analysis

Bioinformatic analysis was performed using Cell Ranger software (Version 1.3.1, 10x Genomics, Pleasanton, CA, USA), the Seurat R package (Satija et al., 2015) and custom software in R and Python, including the *polyseq* Python package developed here. Software to train classifiers using neural networks was built with TensorFlow. The *polyseq* package as well as jupyter notebooks containing code used for analysis in the study are available on GitHub (https://github.com/jwittenbach/polyseq).

Briefly, Cell Ranger was used to perform demultiplexing, alignment, filtering, and counting of barcodes and UMIs, with the output being a cell-by-genes matrix of counts. To further ensure that only high-quality cells were retained, any cell that registered counts in a unique number of genes below a baseline threshold was removed. To reduce the dimensionality of the data for computational tractability, any gene that was not expressed in a baseline number of cells was also dropped.

To account for the fact that raw counts tended to span many orders of magnitude (∼ 10^0^-10^5^), counts were transformed via log(counts + 1). To control for cell size and sequencing depth, the sum of the (log-transformed) counts within each cell used as a regressor for a linear regression model to predict the (log-transformed) counts for each gene (one linear regression model per gene, with each cell being a sample). “Gene expression levels” were then quantified as the z-scored residuals from the fitted models (i.e. standard deviations above/below the predicted log-transformed counts for a particular gene across all cells).

Next, to further reduce the dimensionality of the data in preparation for downstream clustering and embedding operations (both of which have computational costs that scale poorly with the dimensionality of the feature space), principal component analysis was performed with cells as samples and gene expression levels as features. The top K principal components (PCs) were retained as features for downstream analyses. For the lager cell atlas dataset, K was chosen to retain a desired percentage of the total variance. For the smaller cell state datasets, K was chosen automatically via a shuffle test – on each shuffle, gene expression levels for each gene were randomly permuted across all cells and the percent variance explained by the top PC was recorded; the 95^th^ percentile of this value across all shuffles was then used as a threshold to determine the cutoff point for keeping PCs with respect to percent variance explained by a particular PC.

Based on these top PCs, cells were clustered using the Louvain-Jaccard graph-based clustering approach. Briefly, the k-nearest neighbor graph between cells was is computed. Edge weights are then determined using the Jaccard index, which measures the fraction of shared neighbors between any two nodes. Finally, the Louvain community detection algorithm is applied to this graph to partition the nodes into clusters; this algorithm seeks to optimize weight of connections with each cluster relative to those between clusters.

In order to visualize the results of the analysis, the PC features were also used to perform a nonlinear embedding into two dimensions. This was performed via either the t-SNE or the UMAP algorithm.

Once cluster identities were determined, the original gene expression level data was to determine important genes for defining each cluster. For each cluster, gene expression levels were used as features, and a binary indicator of whether or not a cell came from the cluster in question was used as a target. This data was then used to fit a linear classifier (viz, a support vector classifier) to separate in-cluster cells from the rest of the population. The unit normal vector from the linear classifier was then extracted and the components used to rank order genes in terms of importance for defining that cluster. This same technique was also used to find important genes for groups defined by methods other than clustering.

#### Imaging analysis

To quantify the number of vGlut and AstC mRNAs in cells contain both vGlut and AstC, we first manually segmented cells from BB-SIM z-stacks that have both vGlut and AstC FISH signals using the Fiji plugin TrakEM2 (Schindelin et al., 2012; Meissner et al., 2019). After identifying the individual fluorescent spots in segmented cells used a previously described Matlab algorithm (Lionnet et al., 2011), we calculate the number of mRNAs per cell. Reconstructed images were generated using Matlab code that draws spots centered on each of the detected spots positions (Lionnet et al., 2011).

### DATA AND CODE AVAILABILITY

All code and documentation for *polyseq* is open source and freely available on github (https://github.com/jwittenbach/polyseq). Jupyter notebooks used for analysis are available upon request. The scRNA-seq data has been deposited in GEO and is accessible under the accession code GEO: GSE135810.

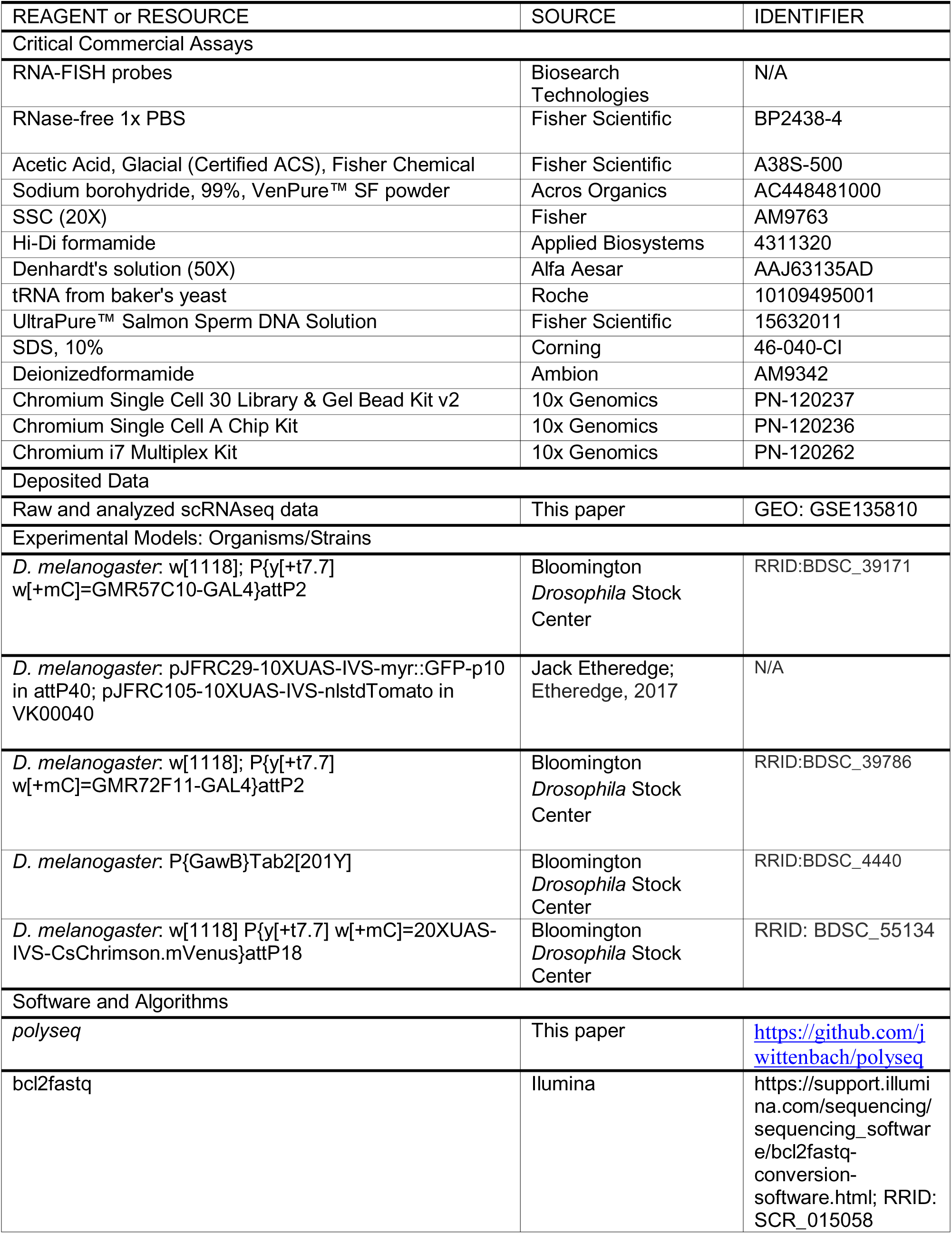
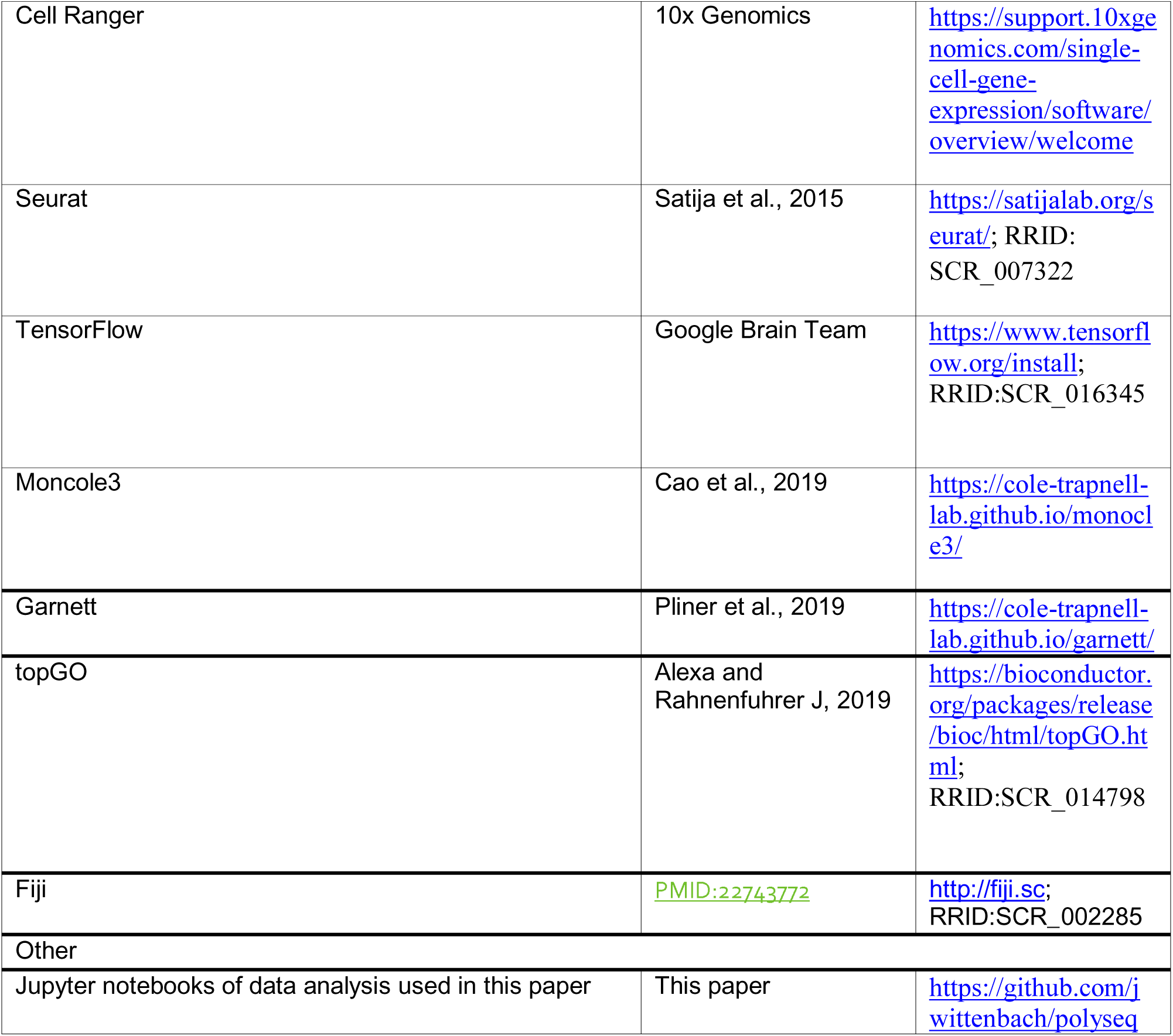

## Supplemental Information titles and legends

**Figure S1.**
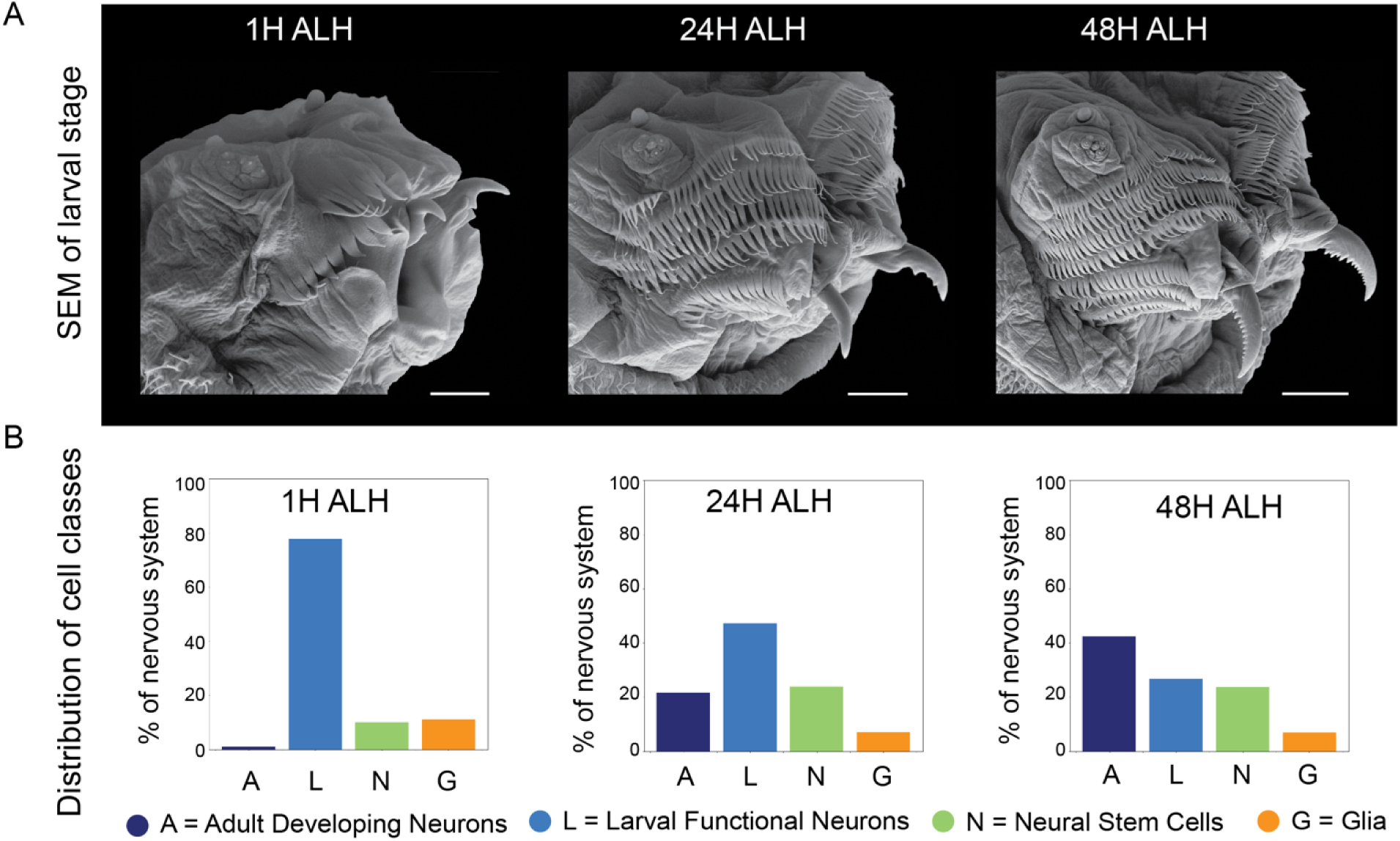
Single cell transcriptomic atlas of the larval nervous system across developmental stages. A. Scanning electron microscopy (SEM) image of larval anatomy during development. Three developmental stages were sequenced separately to build age specific atlases (Scale bar = 10 m for 1H, 20 m for 24H and 40 m for 48 H). Larval size and elaboration of mouth hooks continues during this developmental period. B. Distribution of cell classes. Just after hatching (1 hour after larval hatching (ALH)), there is one primary central cluster of neurons. Neuron classes are recognizable, including cholinergic neurons, motor neurons, astrocyte-like glia, and neuroblasts. At 24 hours ALH multiple main groups of neurons are recognizable with markers consistent with an increase in neuroblasts and ganglion mother cells (GMCs). In addition, there are large groups of neurons, here labeled “developing adult neurons” which have neuron markers but few or no genes expressed for synaptic transmitters and receptors. This is consistent with the large burst of neurons born at this point in development which lay dormant until adult life. At 48 hours ALH the functional larval and developing adult populations are still identifiable.

**Fig S2.**
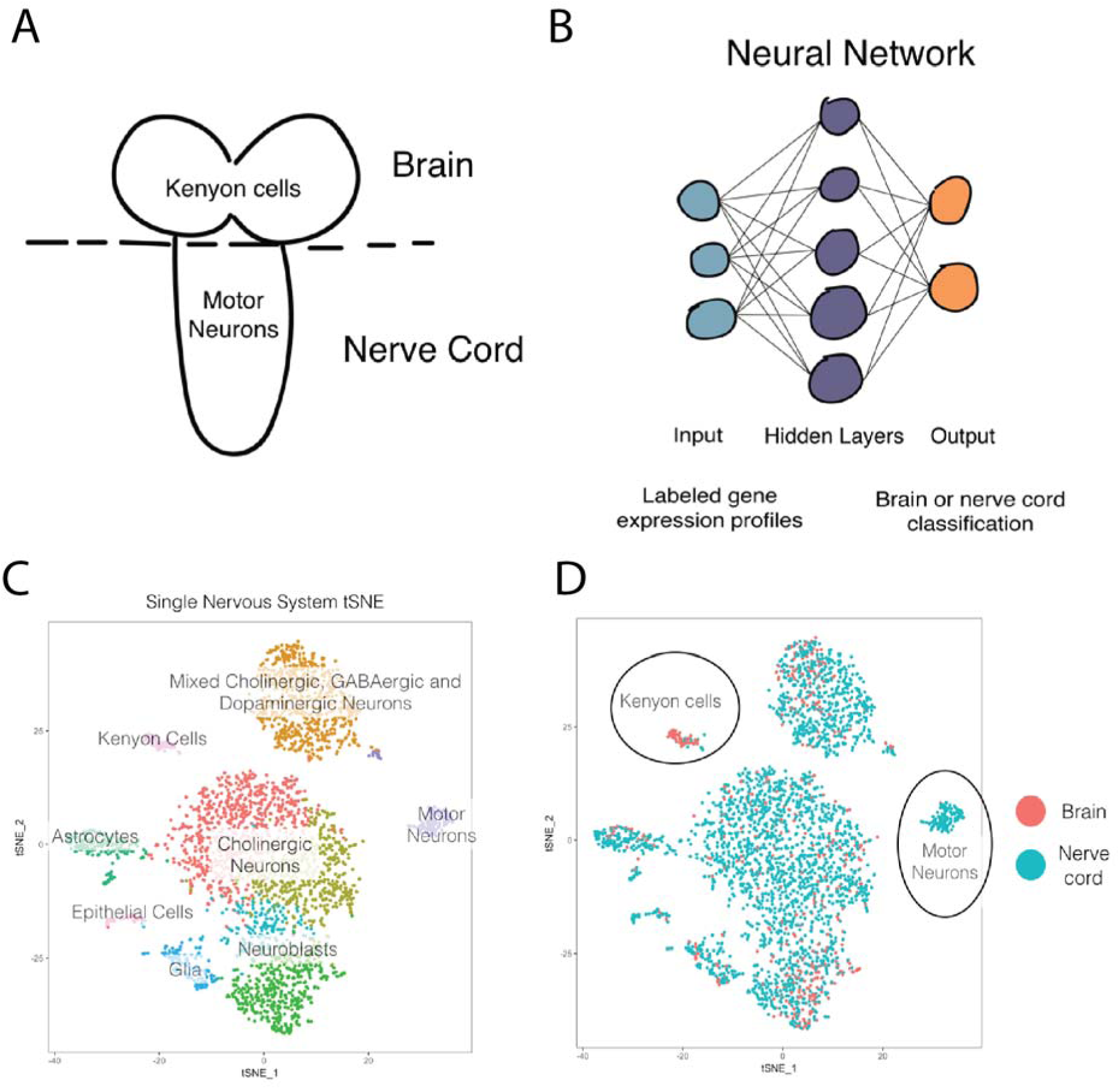
Machine learning separates brain and VNC neurons. We dissected the brain from the nerve cord and sequenced the RNA from each population separately. This provided ground truth labels which we could then feed into (B) a neural network to train a classifier to predict spatial origin from the brain of the VNC. C. We then used our classifier (built on separate data) to predict the brain or VNC origin of cells from a nervous system that was sequenced in one sample (no brian or VNC dissection). D. The classifier correctly separated brain from VNC cells in an intact sample. Most clusters were appropriately mixed brain and VNC cells; however, one pure VNC population (motor neurons) and one pure brain population (Kenyon cells) were appropriately labeled by our machine learning classifier.

**Figure S3.**
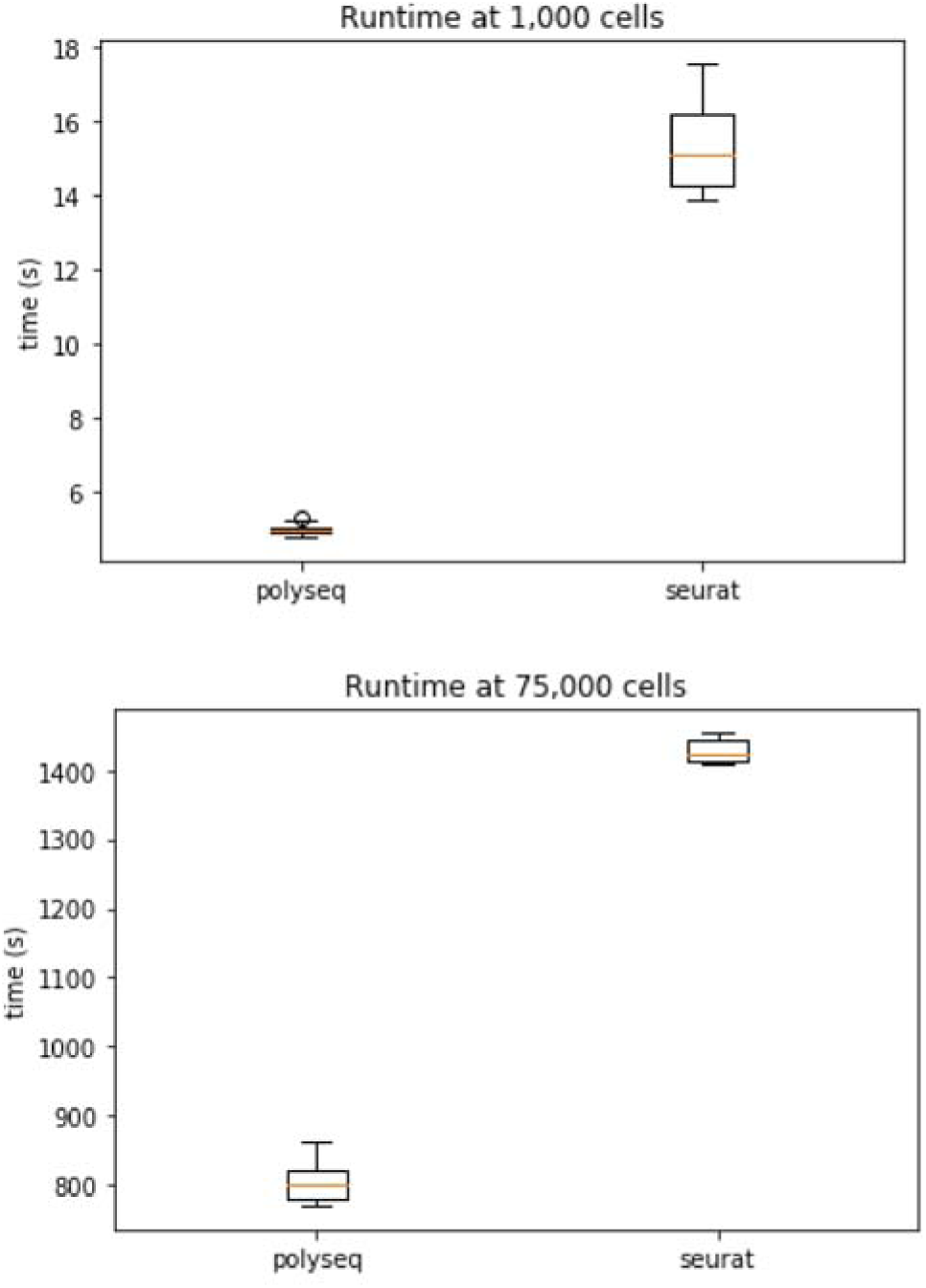
Runtime for a typical pipeline to analyze single cell data in *Seurat* (R) versus *polyseq* (python). Equivalent analysis pipelines were built in *Seurat* and *polyseq* to analyze single cell RNAseq data for datasets of 1,000 cells and 75,000 cells on a single laptop running Mac OS with 16GB RAM and a 2.6 GHz Intel Core i7. Polyseq outperformed Seurat for both small and medium datasets. Jupyter notebooks used for testing that contain code for Seurat and polyseq is available upon request.

**Figure S4.**
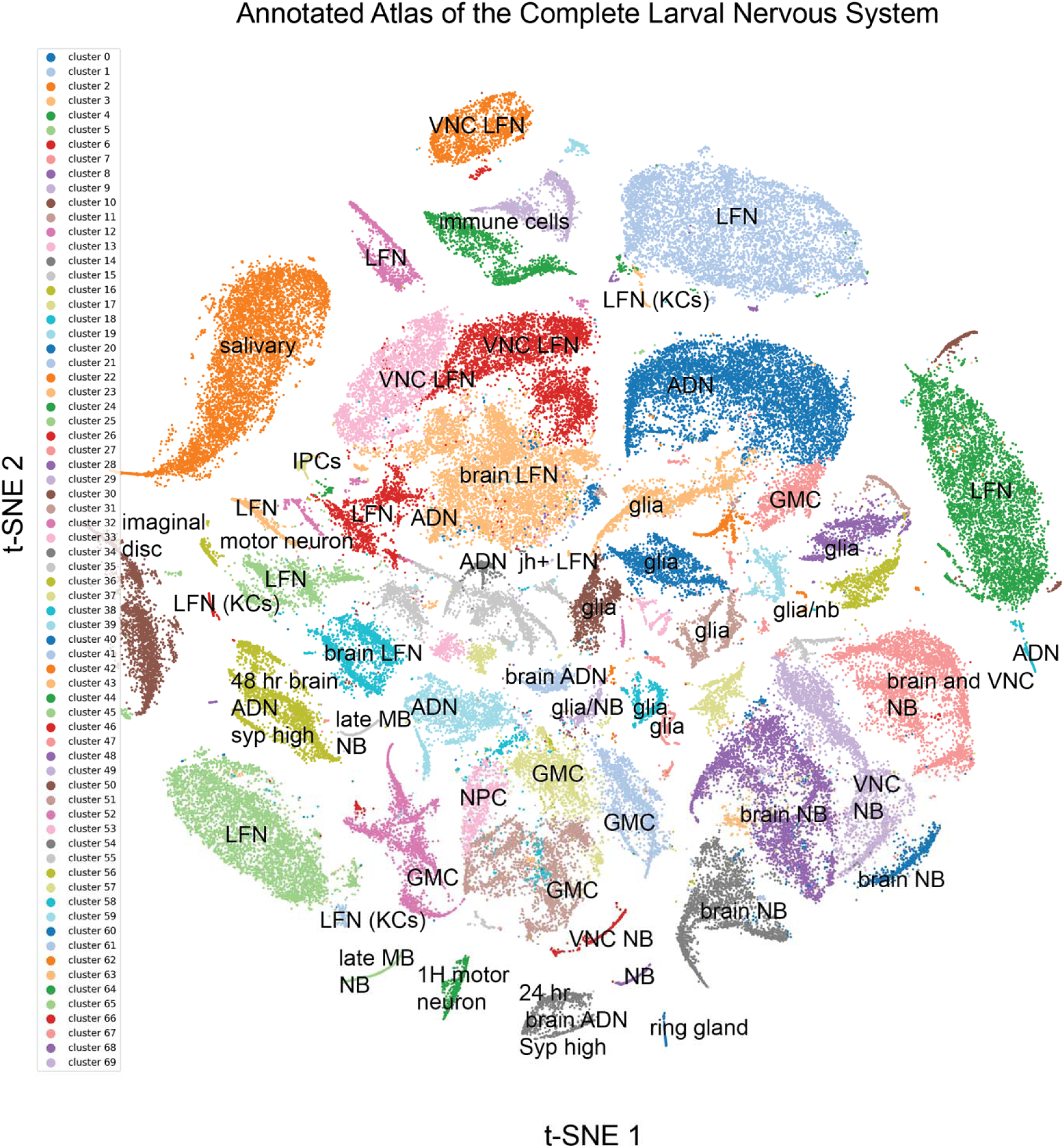
Labeled atlas of the complete larval nervous system. The atlas split into 70 clusters, which are labeled by color and class or more specific name. The top genes which define each cluster, along with the mean expression within the cluster, the expression outside the cluster, and the p-value can be found in xTable S1.

**Figure S4.**
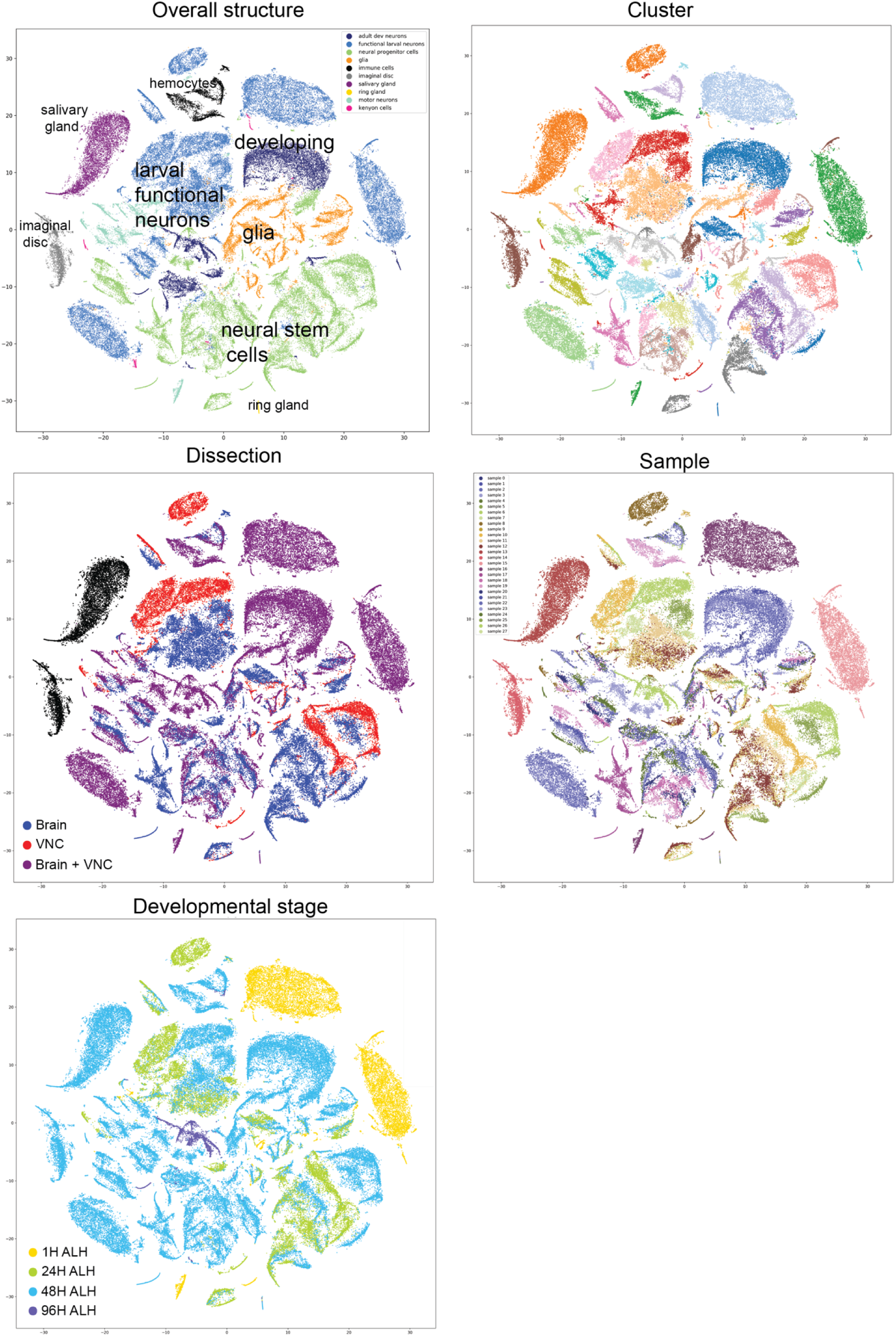
Complete single-cell transcriptomic atlas of the larval central nervous system colored by structure, cluster, dissection, sample, and developmental stage. The complete atlas, described in Figures 4 and S4, colored by key characteristics of the dataset. The first provides an overall structure of the cell classes and where they are found in t-SNE space. The cluster coloring provides information about location for each of the 70 clusters. The dissection splits the data into how the data was collected – whether the sample contained only brain, only VNC, or both. The sample t-SNE provides information about individual 10X genomics samples. The developmental stage provides information about the age of the larvae at collection.

**Figure S6.**
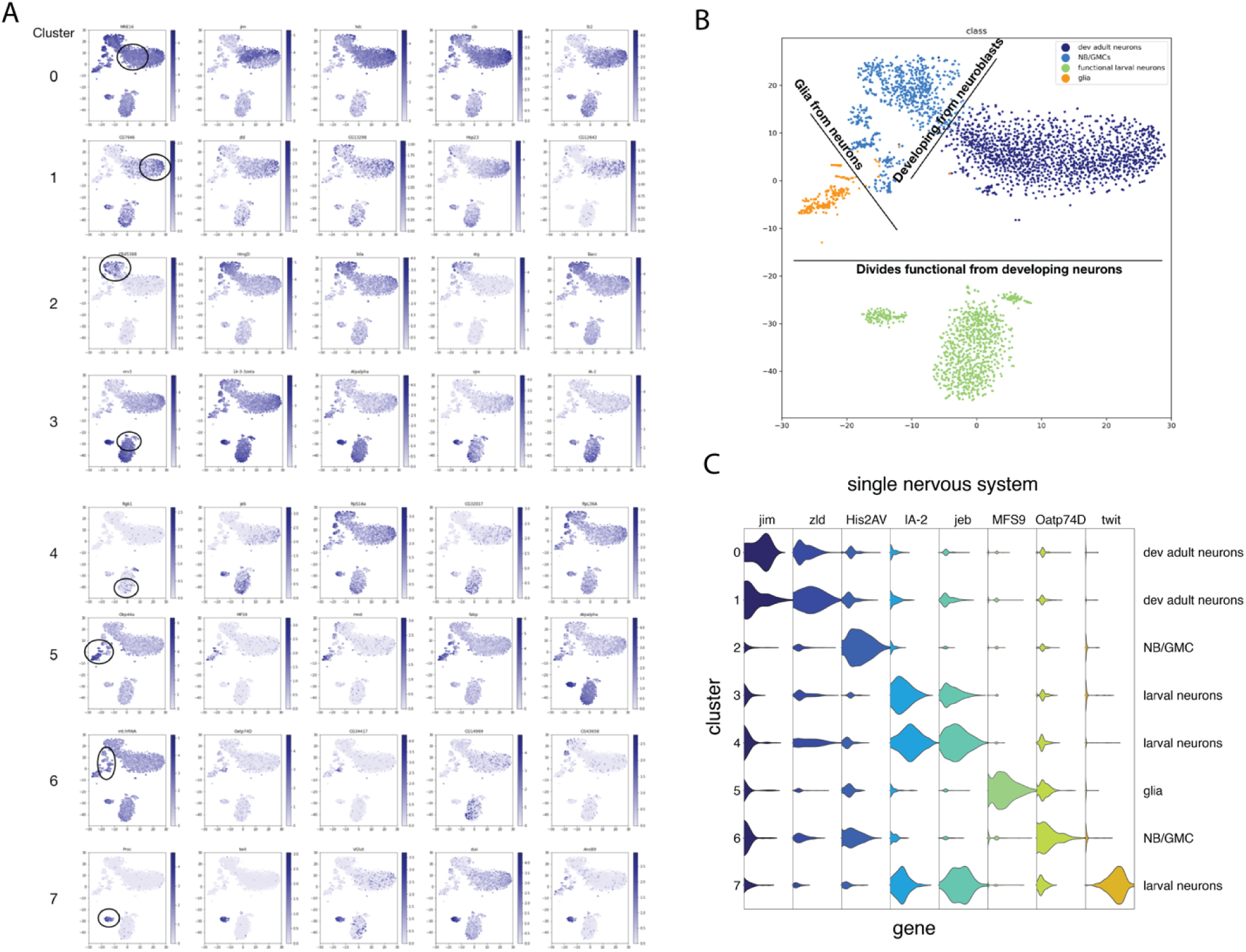
Complete nervous system atlas from an individual animal. A. t-SNE of the complete single nervous system with cell clusters colored by gene expression of the top genes in each cluster. For each cluster in A there is a combination of genes which separate the clusters into recognizable molecular cell types and cell classes. B. Lines can be drawn in the t-SNE space that separates each of the cell classes we define here (adult developing neurons, functional larval neurons, neural stem cells, and glia). C. Violin plot of characteristic genes which separate each of the 8 top level clusters.

**Figure S7.**
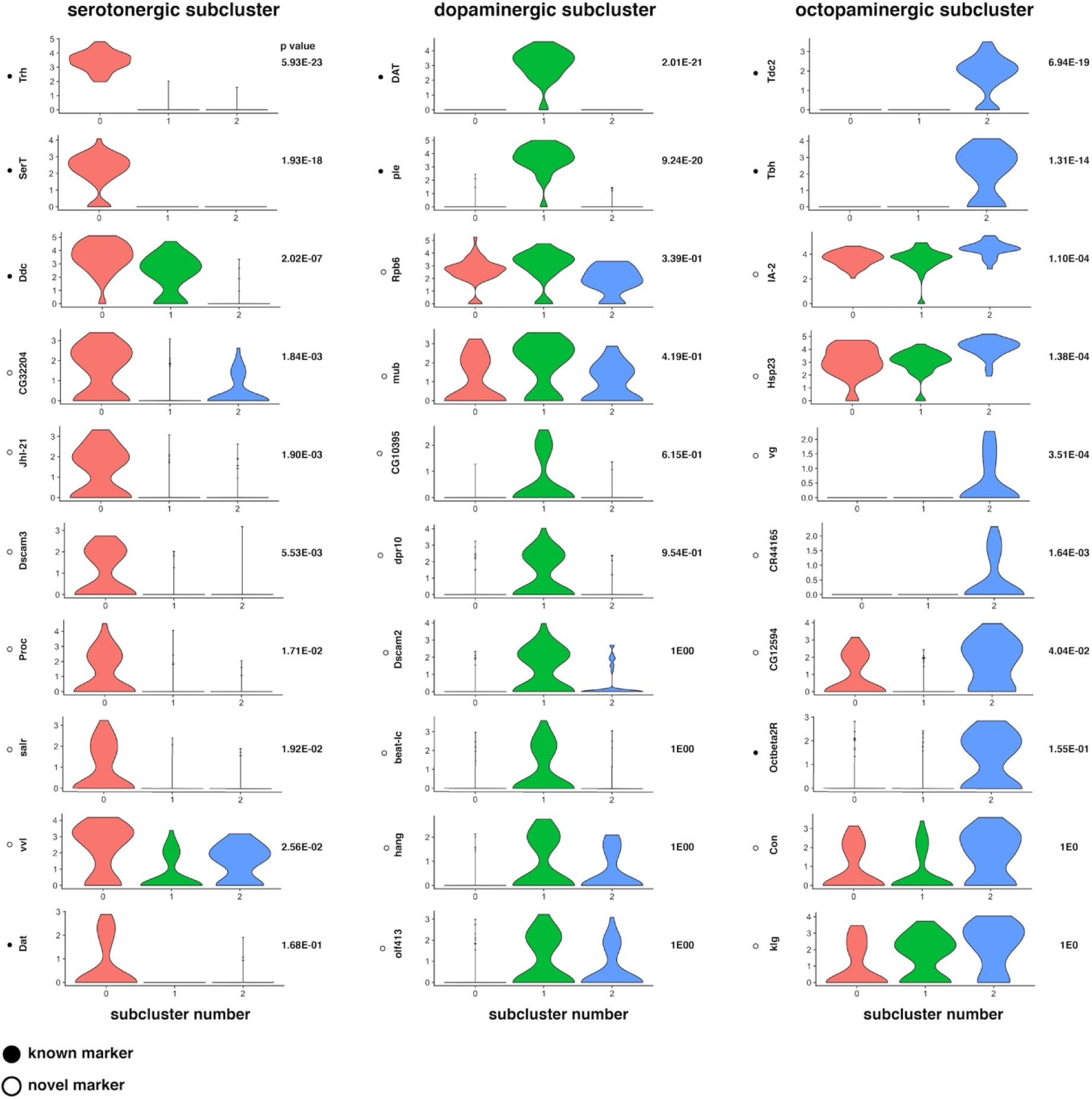
Unsupervised clustering separates recognizable serotonergic, dopaminergic, octopaminergic neurons into subclusters with known validated markers and novel markers. A cluster of cells was discovered using unsupervised machine learning techniques with markers indicative of dopaminergic cells. This cluster was separated and clustered once more, revealing three separate clusters with gene markers indicating a serotonergic subcluster, a dopaminergic subcluster, and an octopaminergic subcluster. The top genes that separate these clusters from one another were then computed and violin plots were generated. Known markers were the top genes for each subcluster and gave them recognizable identity – for example, tryptophan hydroxylase (Trh), which is used to synthesize serotonin from tryptophan, and serotonin transporter (SerT) were the top genes for the first subcluster, which could then be appropriately labeled containing serotonergic cells. In addition to known gene markers, new makers were also discovered for each group of cells.

**Figure S8:**
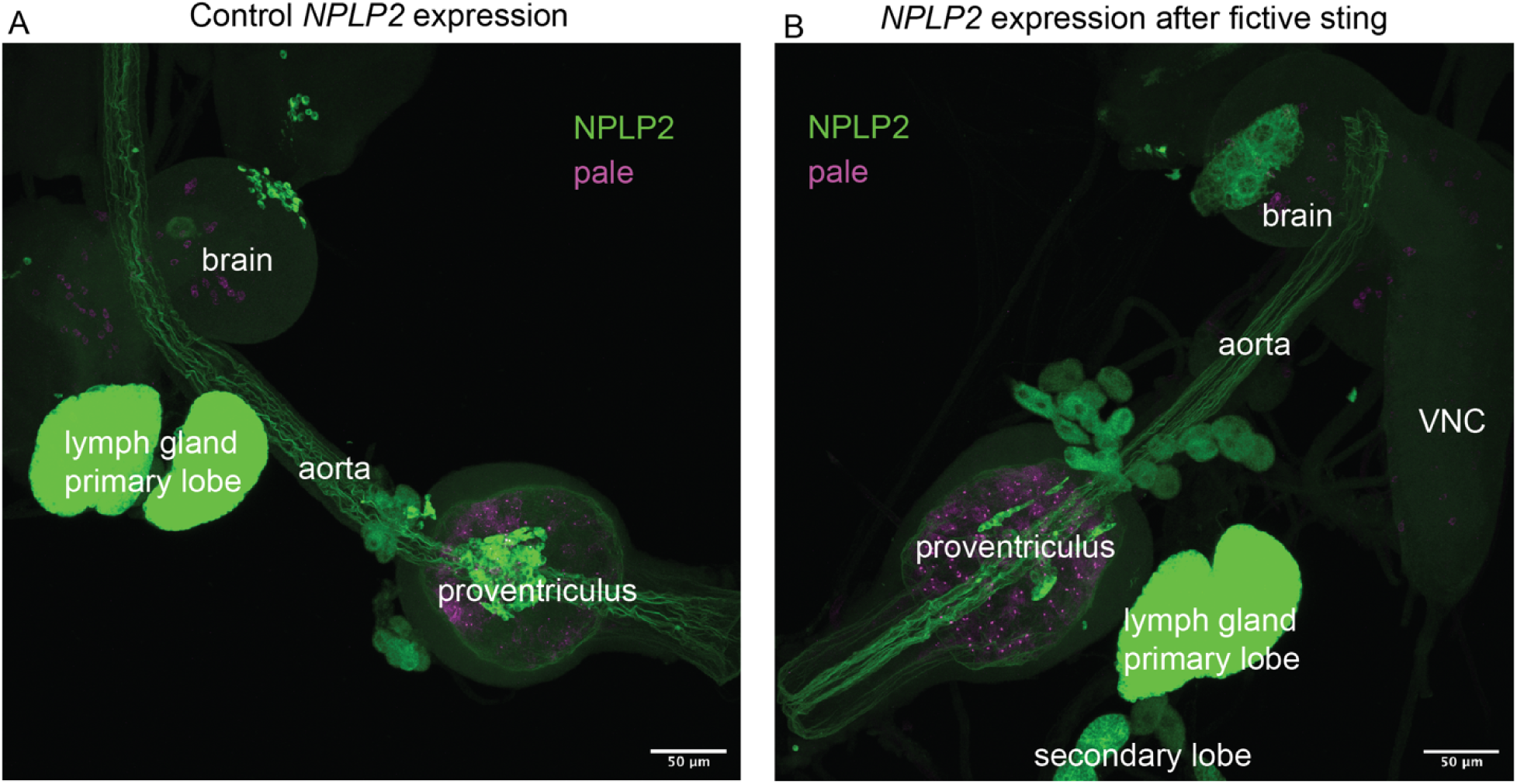
N*P*LP2 expression is increased in the proventriculus and the brain after fictive sting. An RNA-FISH probe was designed for *NPLP2* and *pale* (*ple*) to investigate the strong increased signal of *NPLP2* following the fictive sting. In the brain lobes, the size of the NPLP2-positive cells is much larger. In the proventriculus, where immune cells emerge, there appear to be more NPLP2 positive cells emerging in the case after fictive sting, suggesting an immune reaction releated to the activation of neurons alone.

**Table S1. Atlas of cell types in the complete larval nervous system.**

The top genes for the 70 cell type clusters are provided with the mean expression inside the cluster, mean expression outside the cluster, and p-value. These clusters correspond to Figure 4 and Supplementary Figure S4.

**Table S2. Gene modules that characterize neural precursor cells over development.**

Differential gene expression was used to compute gene modules in the Monocle3 R package. The modules characterize early, intermediate, and late neural precursor cell populations (Figure 3).

**Table S3. Fictive sting full nervous system atlas.**

The top genes for the 14 cell type clusters obtained from animals that were fictively stung and controls. Tables include the top genes for each cluster, the mean expression inside the cluster, the mean expression outside the cluster, and the p-value. These clusters correspond to Figure 7E-F.

**Table S4. KC overactivation full nervous system atlas.**

The top genes for the 14 cell type clusters obtained from animals that had KCs repeatedly activated and controls. Tables include the top genes for each cluster, the mean expression inside the cluster, the mean expression outside the cluster, and the p-value. These clusters correspond to Figure 7G-I.

**Table S5: RNA-FISH probe sequences.**

RNA sequeces of probes built to label *AstC-R2*, *Hug*, and *NPLP2* transcripts.

**Movie S1. Z-stacks through RNA-FISH of AstC in insulin-producing cells.**

Z stacks are shown through the confocal stack of a brain in which the IPCs are labeled with a fluorescent HaloTag ligand (Magenta), and AstC-R2 mRNA is detected by FISH (green), bar 10µm. This movie corresponds to FIgure 5 in the main text.

**Movie S2. Single-molecule imaging to correlate between single-molecule FISH and scRNAseq.**

Cells were identified with coexpression of AstC (green) and vGlut (magenta) in the whole brain. This movie corresponds to Figure 6 in the main text and shows a view through z-stacks of BB-SIM images of AstC and vGlut mRNA FISH channels.

